# Discovery of several thousand highly diverse circular DNA viruses

**DOI:** 10.1101/555375

**Authors:** Michael J. Tisza, Diana V. Pastrana, Nicole L. Welch, Brittany Stewart, Alberto Peretti, Gabriel J. Starrett, Yuk-Ying S. Pang, Siddharth R. Krishnamurthy, Patricia A. Pesavento, David H. McDermott, Philip M. Murphy, Jessica L. Whited, Bess Miller, Jason M. Brenchley, Stephan P. Rosshart, Barbara Rehermann, John Doorbar, Blake A. Ta’ala, Olga Pletnikova, Juan Troncoso, Susan M. Resnick, Ben Bolduc, Matthew B. Sullivan, Arvind Varsani, Anca M. Segall, Christopher B. Buck

## Abstract

Although it is suspected that there are millions of distinct viral species, fewer than 9,000 are catalogued in GenBank’s RefSeq database. We selectively enriched for and amplified the genomes of circular DNA viruses in over 70 animal samples, ranging from cultured soil nematodes to human tissue specimens. A bioinformatics pipeline, Cenote-Taker, was developed to automatically annotate over 2,500 circular genomes in a GenBank-compliant format. The new genomes belong to dozens of established and emerging viral families. Some appear to be the result of previously undescribed recombination events between ssDNA viruses and ssRNA viruses. In addition, hundreds of circular DNA elements that do not encode any discernable similarities to previously characterized sequences were identified. To characterize these “dark matter” sequences, we used an artificial neural network to identify candidate viral capsid proteins, several of which formed virus-like particles when expressed in culture. These data further the understanding of viral sequence diversity and allow for high throughput documentation of the virosphere.

## Introduction

There has been a rush to utilize the massive parallel sequencing approaches to better understand the complex microbial communities associated with humans and other animals. Although the bacterial populations in these surveys have become increasingly recognizable (Lloyd-Price et al., 2017), a substantial fraction of the reads and *de novo* assembled contigs in many metagenomics efforts are binned as genetic “dark matter,” with no recognizable similarity to characterized sequences (Krishnamurthy and Wang, 2017, Oh et al., 2014). Some of this dark matter undoubtedly consists of viral sequences, which have remained poorly characterized due to their enormous diversity (Simmonds et al., 2017, Paez-Espino et al., 2016, Emerson et al., 2018). Recent efforts have shown that our understanding of viral diversity, even of viruses known to directly infect humans, has been incomplete (Pastrana et al., 2018, Turnbaugh et al., 2007, Gilbert et al., 2010). To increase the power of future studies seeking to more comprehensively catalog the virome and find additional associations between viruses and disease, reference genomes for all clades of the virosphere need be identified, annotated, and made publicly accessible.

Virus discovery has typically proven to be more difficult than discovery of cellular organisms. Whereas all known cellular organisms encode conserved sequences (such as ribosomal RNA genes) that can readily be identified through sequence analysis, viruses, as a whole, do not have any universally conserved sequence components (O’Leary et al., 2016, Brister et al., 2015, Sullivan, 2015, Rohwer and Edwards, 2002). Nevertheless, some success has been achieved in RNA virus discovery by probing for the conserved sequences of their distinctive RNA-dependent RNA polymerase or reverse transcriptase genes in metatranscriptomic data (Shi et al., 2016). Also, many bacteriophages of the order *Caudovirales*, such as the families *Siphoviridae*, *Podoviridae*, and *Myoviridae*, have been reported in high numbers due to their and their hosts’ culturability and their detectability using viral plaque assays (Pope et al., 2015, Grose and Casjens, 2014, Grose et al., 2014). The relatively abundant representation of these families in databases has allowed new variants to be recognized by high-throughput virus classification tools like VirSorter (Roux et al., 2015, Gregory et al., 2019, Roux et al., 2019b). In contrast, many small DNA viruses are not easily cultured (Bedell et al., 1991), use diverse genome replication strategies, and typically lack DNA polymerase genes such as those in large DNA viruses (Koonin et al., 2015). An additional challenge is that small DNA viruses with segmented genomes may have segments that do not encode recognizable homologs of known viral genes. Therefore, small DNA viruses are more sparsely represented in reference databases. However, some groups have been successful in discovery of small DNA genomes in a wide range of viromes (Blinkova et al., 2010, Pastrana et al., 2018, Dayaram et al., 2015, Dayaram et al., 2016, Labonte and Suttle, 2013, Rosario et al., 2018, Victoria et al., 2009).

Despite the apparent challenges in detecting small DNA viruses, many have physical properties that can be leveraged to facilitate their discovery. In contrast to the nuclear genomes of animals, many DNA virus genomes have circular topology, which allows selective enrichment through rolling circle amplification (RCA) methods (Kim et al., 2008). Further, the unique ability of viral capsids to protect nucleic acids from nuclease digestion and to mediate the migration of the viral genome through ultracentrifugation gradients or size exclusion columns allows physical isolation of viral genomes.

The current study grew out of an effort to find papillomaviruses (small circular DNA viruses) in humans and economically important or evolutionarily informative animals (Pastrana et al., 2018, Peretti et al., 2015). The sampling included several types of animals that might serve as laboratory models (e.g., mice, fruit flies, soil nematodes). A number of papillomaviruses were detected among a vastly larger set of circular DNA sequences that were not easily identifiable in standard BLASTN searches. The goal of the present study is to catalog and annotate the circular DNA virome from these animal tissues to understand the diversity and evolution of viral sequences. We developed a comprehensive bioinformatics pipeline, Cenote-Taker, to classify and annotate over 2,500 candidate viral genomes and generate GenBank-compliant output files. Cenote-Taker is available for free public use with a graphical user interface at http://www.cyverse.org/discovery-environment.

## Results

### Virion enrichment, genome sequencing, and annotation

We have previously developed methods for discovery of new polyomavirus and papillomavirus species in skin swabs and complex tissue specimens (Peretti et al., 2015). Nuclease-resistant DNA from purified virions was amplified by random-primed rolling circle amplification (RCA) and subjected to deep-sequencing. Reads were *de novo* assembled into contigs and analyzed with a bioinformatics pipeline, Cenote-Taker (a portmanteau of *cenote*, a naturally occurring circular water pool, and *note-taker*), to identify and annotate *de novo* assembled contigs with terminal direct repeats consistent with circular DNA molecules. In this pipeline, putative circular sequences were queried against GenBank’s nucleotide database using BLASTN to remove circles with extensive nucleotide identity (>90% across any 500 bp window) to known sequences. Sequences with >90% identity to previously reported viral sequences represented less than 1.5% of circular contigs and are not included in further analysis. Open reading frames (ORFs) from remaining unidentified circular DNA sequences >240 nucleotides (nt) in length were translated and used for RPS-BLAST queries of GenBank’s Conserved Domain Database (CDD). ORFs that did not yield E values better than 1×10^−4^ in RPS-BLAST were subjected to BLASTP searches of viral sequences in GenBank’s nr database (Altschul et al., 1990, Marchler-Bauer and Bryant, 2004, Marchler-Bauer et al., 2015). For ORFs that were not confidently identified in BLAST searches, HHBlits (Remmert et al., 2011) was used to search the CDD, Pfam (El-Gebali et al., 2019), Uniprot (UniProt, 2019), Scop (Chandonia et al., 2019), and PDB (Burley et al., 2017) databases. The results were used to annotate and name each sequence in a human-readable genome map as well as a format suitable for submission to GenBank. After checking the Cenote-Taker output of each genome, minor revisions were made, as needed, and files were submitted to GenBank (BioProject Accessions PRJNA393166 and PRJNA396064). All annotations meet or exceed recently proposed standards for uncultivated virus genomes (Roux et al., 2019a).

### Discovery of 2514 DNA viruses in animal metagenomes

Of the novel circular sequences detected in the survey, 1844 encode genes with similarity to proteins of ssDNA viruses and 55 encode genes with similarity to dsDNA viral proteins (Fig. 1A). The large majority of genomes from this study are highly divergent from RefSeq entries (Fig. S1). We discovered 868 genomes that had similarity to unclassified eukaryotic viruses known as circular replication associated protein (Rep)-encoding single-stranded DNA (CRESS) viruses. The group is defined by the presence of a characteristic rolling circle endonuclease/superfamily 3 helicase gene (Rep) (Zhao et al., 2019, Kazlauskas et al., 2019), but has not been assigned to families by the ICTV or RefSeq. We estimate that 199 non-redundant unclassified CRESS virus genomes had been previously deposited in GenBank, and 85 are listed in RefSeq (Fig. 1B). Also abundant was the viral family *Microviridae*, a class of small bacteriophages, with 670 complete genomes. This represents a substantial expansion beyond the 459 non-redundant microvirus genomes previously listed in GenBank (of which 44 were listed in the RefSeq database). Other genomes that were uncovered represent *Anelloviridae* (n=170), *Inoviridae* (n=70), *Genomoviridae* (n=58), *Siphoviridae* (n=18), unclassified phage (n=14), *Podoviridae* (n=10), *Myoviridae* (n=7) unclassified virus (n=6), *Papillomaviridae* (n=4), *Circoviridae* (n=3), unclassified *Caudovirales* (n=3), *Bacilladnaviridae* (n=2), *Smacoviridae* (n=2), and *CrAssphage* (n=2) (Fig. 1B).

**Figure 1:**
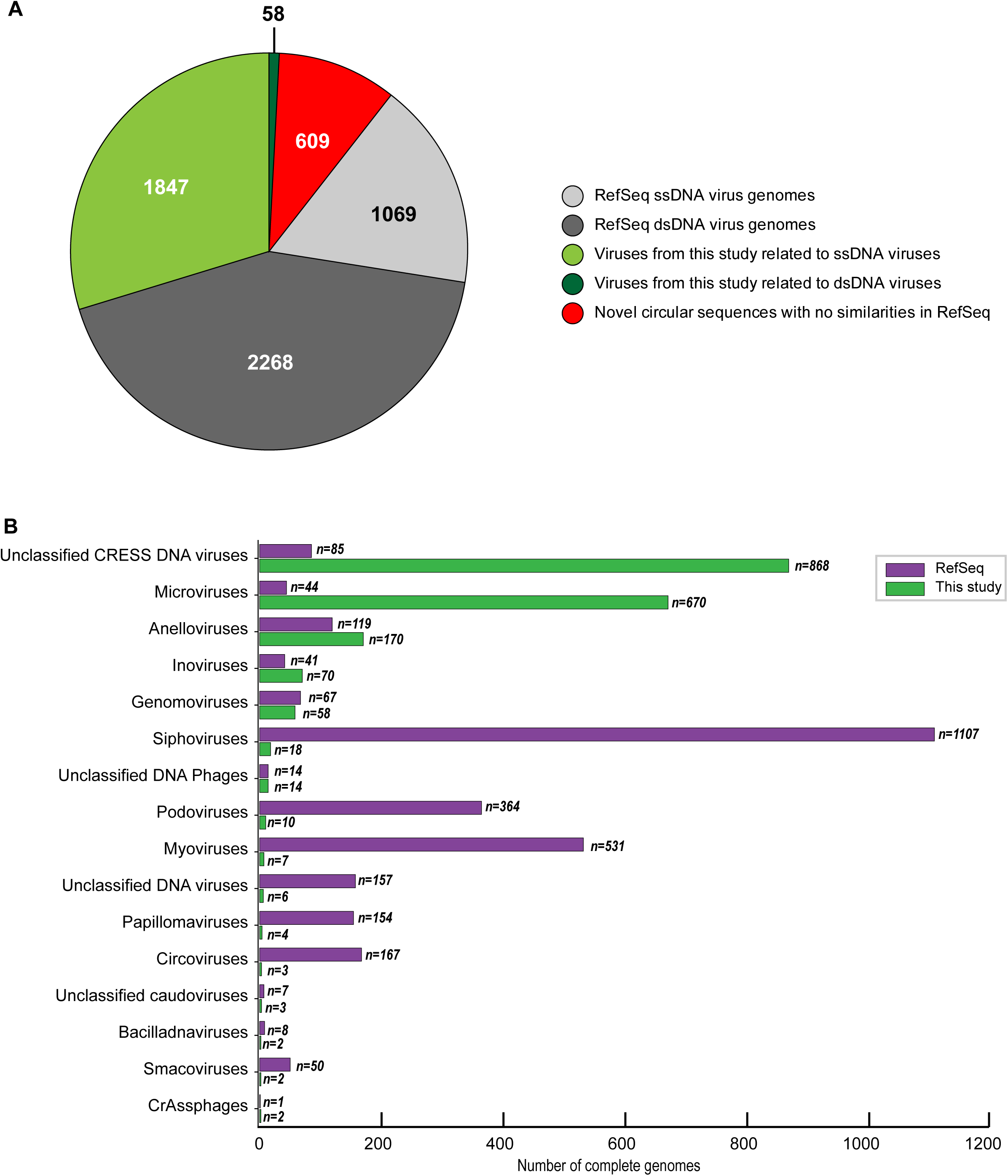
Novel viruses associated with animal samples. Gross characterization of viruses discovered in this project compared to NCBI RefSeq virus database entries. (A) Pie chart representing the number of viral genomes in broad categories. (B) Bar graph showing the number of new representatives of known viral families or unclassified groups. Note that genomes in this study were assigned taxonomy based on at least one region with a BLASTX hit with an E value <1×10^−5^, suggesting commonality with a known viral family. Some genomes may ultimately be characterized as being basal to the assigned family.

It is difficult to assign a host to most of the viruses from this study due to their divergence from known viral sequences. However, we searched the CRISPR database at (https://crispr.i2bc.paris-saclay.fr/crispr/BLAST/CRISPRsBlast.php), and three viruses had exact matches to CRISPR spacers in bacterial genomes (Siphoviridae sp. ctcj11:Shewanella sp. W3-18-1, Inoviridae sp. ctce6:Shewanella baltica OS195, Microviridae sp. ctbe523:Paludibacter propionicigenes WB4) and one virus had an exact match to the CRISPR spacer of an archaeon (Caudovirales sp. cthg227:Methanobrevibacter sp. AbM4), implying that these organisms are infected by these viruses. Further, the 142 anelloviruses found in human blood samples (Table S1) are almost certain to be bona fide human viruses based on their relatedness to known human anelloviruses.

In addition to circular genomes with recognizable similarity to known viruses, 609 circular contigs appeared to represent elements that lacked discernable similarity to known viruses (Fig. 1A).

The vast majority of the *de novo* assembled circular genomes were <10 kb in length (Fig S2). This is largely due to the fact that large genomes are typically more difficult to *de novo* assemble from short reads. Despite these technical obstacles, our detection of a new tailed bacteriophage with a 419kb genome (Myoviridae sp. isolate ctbc_4, GenBank Accession: MH622943), along with 45 other >10 kb circular sequences (Fig S2), indicates that the methods used for the current work can detect large viral genomes.

There has been a recent renewal of interest in the hypothesis that viruses may be etiologically associated with degenerative brain diseases, such as Alzheimer’s disease (Itzhaki et al., 2016, Eimer et al., 2018). Conflicting literature suggests the possible presence of papillomaviruses in human brain tissue (Coras et al., 2015, Chen et al., 2012). Samples of brain tissue from individuals who died of Alzheimer’s disease (n=6) and other forms of dementia (n=6) were subjected to virion enrichment and deep sequencing. Although complete or partial genomes of known papillomaviruses, Merkel cell polyomavirus, and/or anelloviruses were observed in some samples (Supplementary File S1), no novel complete viral genomes were recovered (Table S1). No viral sequences were detected in a follow-up RNA deep sequencing analysis of the brain samples. It is difficult to know how to interpret these negative data. It is conceivable that the known viral DNA sequences observed in the Optiprep-RCA samples represent virions from blood vessels or environmental sources.

It has recently become apparent that certain nucleic acid extraction reagents are contaminated with viral nucleic acids (Asplund et al., 2019). To ensure we were not merely reporting the sequences of the “reagent virome,” we performed our wet bench and bioinformatic pipeline on three independent replicates of reagent-only samples. We found no evidence of sequences of any viruses reported here or elsewhere. Further, cross-sample comparison of contigs showed that almost no sequences were found in different animal samples, aside from technical replicates. In total, six viral genomes were observed in multiple unrelated samples from at least two sequencing runs (Table S2). It is unclear whether this small minority of genomes (0.24% of the genomes reported in the current study) represent reagent contamination, lab contamination, or actual presence of the sequences in different types of samples.

### Assignment of hallmark genes to networks shows expansion of virus sequence space

Single stranded DNA viruses, in general, have vital genes encoding proteins that mediate genome replication, provide virion structure, and, in some cases, facilitate packaging of viral nucleic acid into the virion. Being structurally conserved, these genes also tend to be important for evolutionary comparisons and can serve as important “hallmark genes” for virus discovery and characterization. However, even structurally conserved proteins sometimes do not have enough sequence conservation as to be amenable to high confidence BLASTP searches. We therefore set out to catalog hallmark ssDNA virus genes based using protein structural prediction. Structures of hallmark genes of exemplar isolates from most established ssDNA virus families have been solved and deposited in publicly available databases, such as PDB (Protein Data Bank) (Burley et al., 2017). Using bioinformatic tools, such as HHpred, one can assign structural matches for a given gene based on the predicted potential folds of a given amino acid sequence. HHpred has been extensively tested and validated for computational structural modelling by the structural biology community (Meier and Soding, 2015, Huang et al., 2014). The method proves especially useful for protein sequences from highly divergent viral genomes that have little similarity to annotated sequences in current databases.

We extracted protein sequences from our dataset and compiled nonredundant proteins from circular ssDNA viruses in GenBank and used them as queries in HHpred searches against the PDB, PFam, and CDD databases. We then grouped structurally identifiable sequences into hallmark gene categories and aligned them pairwise (each sequence was compared to all other sequences) using EFI-EST (Gerlt et al., 2015). The resulting sequence similarity networks (SSNs) were visualized with Cytoscape (Su et al., 2014), with each node representing an predicted protein sequence (Fig. 2–3, Fig. S3). Nodes (sequences) with significant amino acid similarity are connected with lines representing BLAST similarity scores below a threshold E value. Sequence similarity network analyses, it has been proposed (Iranzo et al., 2017), represent relationships between viral sequences better than phylogenetic trees. Further, SSNs have previously been used for viral protein and genome cluster comparison (Bolduc et al., 2017, Lima-Mendez et al., 2008, Lefeuvre et al., 2019, Kazlauskas et al., 2019), and can be used to display related groups of viral genes in two dimensions (Bin Jang et al., 2019). These clusters were also used to guide the construction of meaningful phylogenetic trees (Fig. 2A-B, Fig. S4).

**Figure 2:**
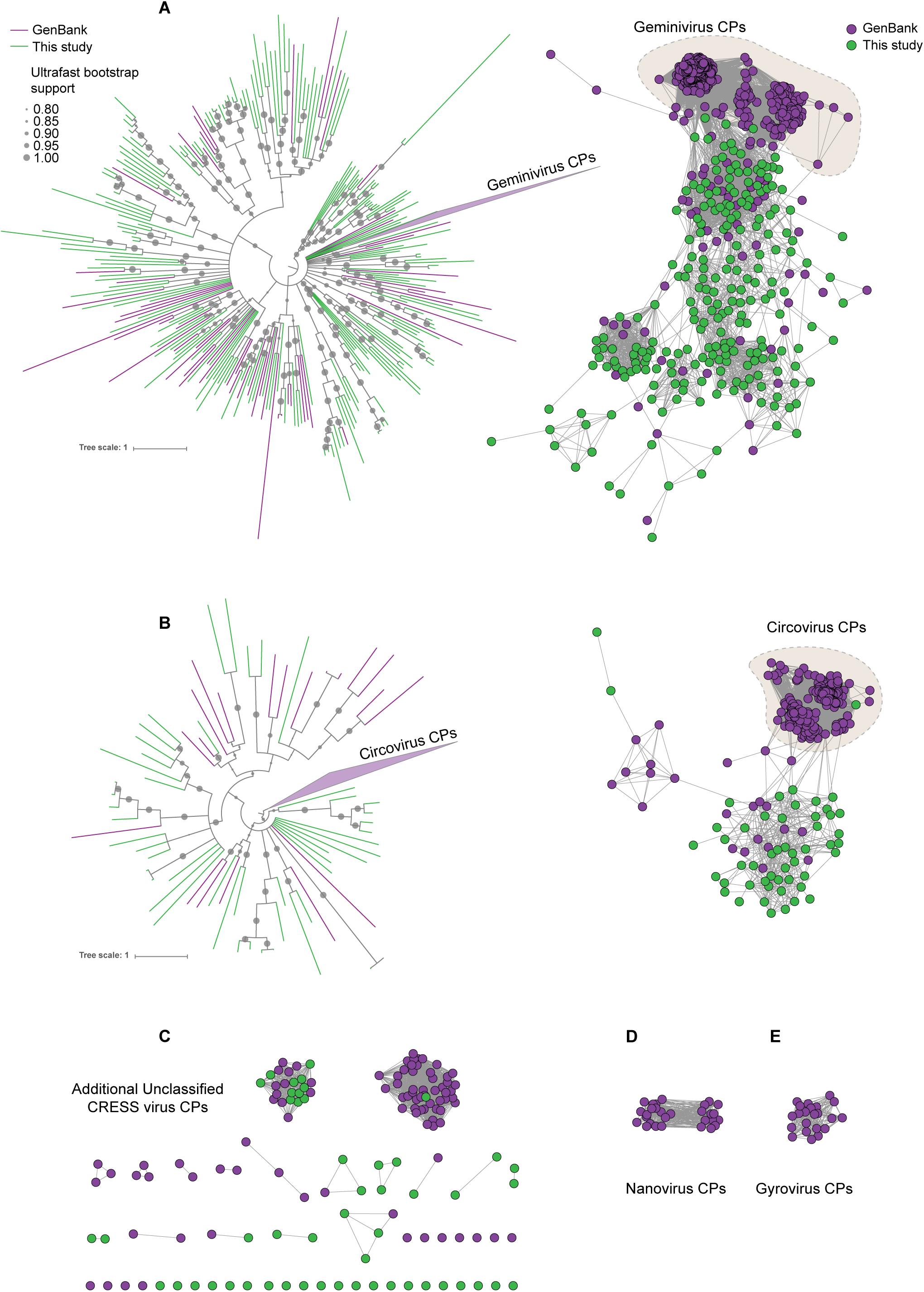
Sequence similarity network analysis of CRESS virus capsid proteins. EFI-EST was used to conduct pairwise alignments of amino acid sequences from this study and GenBank with predicted structural similarity to CRESS virus capsid proteins. The E value cutoff for the analysis was 10^−5^. (A) Cluster consisting of proteins with predicted structural similarity to geminivirus-like capsids and/or STNV-like capsids. The phylogenetic tree was made from all sequences in this cluster. (B) A cluster consisting of sequences with predicted structural similarity to Circovirus capsid proteins. The phylogenetic tree was made from all sequences in this cluster. (C) Assorted clusters and singletons from unclassified CRESS virus proteins that were modelled to be capsids. (D) Nanovirus capsids. (E) Gyrovirus capsids.

**Figure 3:**
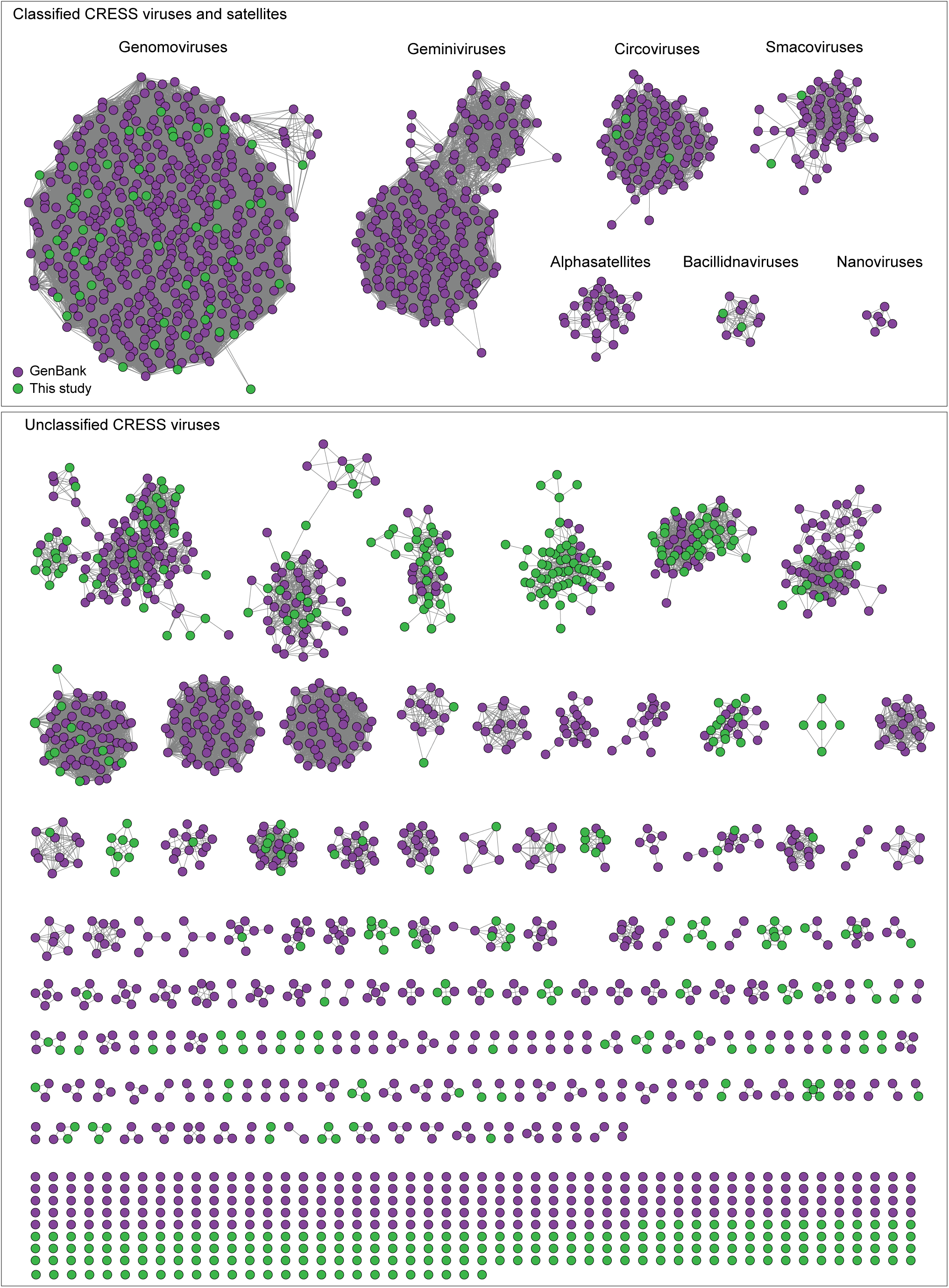
Network analysis of CRESS virus Rep proteins. EFI-EST was used to conduct pairwise alignments of amino acid sequences from this study and GenBank that were structurally modelled to be a rolling-circle replicase (Rep). The analysis used an E value cutoff of 10^−60^ to divide the data into family-level clusters.

In Figure 2, sequences that showed a structural match to a known eukaryotic circular ssDNA virus capsid protein are displayed as a network. This general capsid type features a single beta-jellyroll fold and assembles into T=1 virions of 20-30 nm in diameter. The network shows that sequences from this study expand and link smaller disconnected clusters of sequences found in GenBank entries (Fig. 2A-C). Perhaps more importantly a number of previously unknown clusters were identified, providing insight into highly divergent hallmark sequences and making this capsid sequence space amenable to BLAST searches in GenBank (Fig. 2C). Although the satellite tobacco necrosis virus (STNV) capsid protein encapsidates an RNA molecule, it has previously been noted that its structure is highly similar to the capsid proteins of geminiviruses and other ssDNA viruses (Koonin et al., 2015, Kraberger et al., 2015, Krupovic et al., 2009, Hipp et al., 2017, Bottcher et al., 2004, Zhang et al., 2001) and was included as a model for populating this network.

A similar pattern can be seen in sequence similarity networks for the Rep genes of CRESS viruses (Fig. 3). Rep genes have been the primary sequences used for taxonomy of CRESS viruses (Zhao et al., 2019). In this case, it was determined that a network with alignment cutoffs with E values of 1×10^−60^ could split the data neatly into “family-level” clusters (Fontenele et al., 2019, Kraberger et al., 2019), precisely mirroring ICTV taxonomy of CRESS viruses. Many additional family-level clusters can be discerned from unclassified CRESS viruses. Other eukaryotic and prokaryotic ssDNA virus hallmark gene networks are shown in Supplementary Figure 3. Phylogenetic trees of networks from Figure 3 and Supplemental Figure 3 are displayed in Supplemental Figure 4.

Cytoscape files of sequence similarity networks and phylogenetic trees can be found at https://ccrod.cancer.gov/confluence/display/LCOTF/DarkMatter.

### New classes of large CRESS viruses feature unconventional structural genes

Although no single family of viruses accounts for the majority of genomes in this study, these results expand the knowledge of the vast diversity of CRESS viruses, which appear to be ubiquitous among eukaryotes (Krupovic et al., 2016, Zerbini et al., 2017, Rosario et al., 2017, Varsani and Krupovic, 2018) and are likely to also infect archaea (Díez-Villaseñor and Rodriguez-Valera, 2019, Kazlauskas et al., 2019). Characterized CRESS viruses have small icosahedral virions (20-30 nm in diameter) with a simple T=1 geometry (Khayat et al., 2011). This capsid architecture likely limits genome size, as nearly all previously reported CRESS virus genomes and genome segments are under 3.5 kb. Exceptions to this size rule are bacilladnaviruses, which have 4.5 - 6 kb genomes (Tomaru et al., 2011) and cruciviruses, which have 3.5 - 5.5 kb genomes (Quaiser et al., 2016). Interestingly, the genomes of these larger CRESS viruses encode capsid genes that appear to have been acquired horizontally from RNA viruses (Kazlauskas et al., 2017). In our dataset, eight CRESS-like circular genomes exceed 6 kb in length (Fig. 4C-J). Further, this study’s large CRESS genomes are apparently attributable to several independent acquisitions of capsid genes from other taxa and/or capsid gene duplication events.

**Figure 4:**
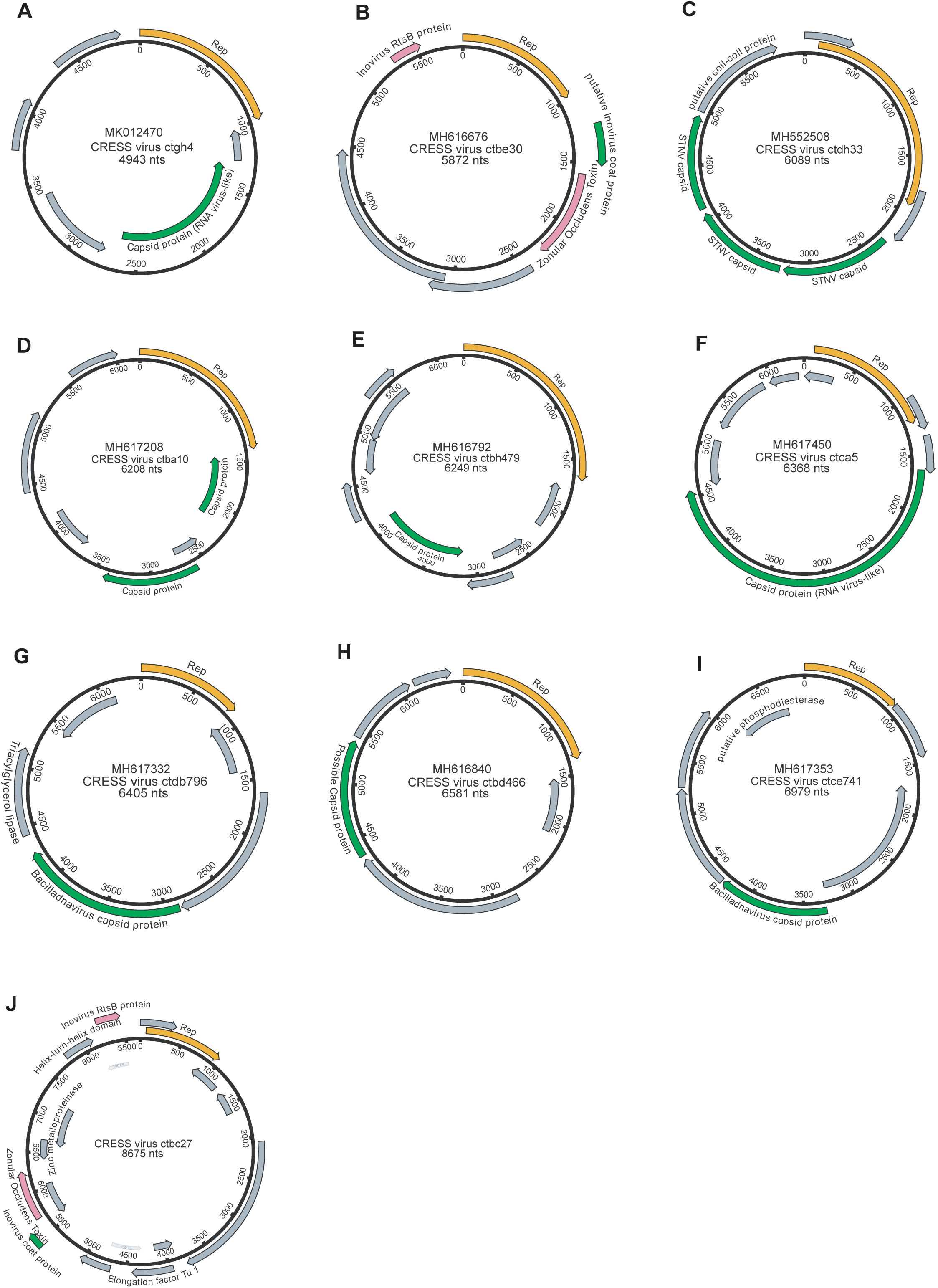
Genome maps of large CRESS virus genomes. Predicted CRESS Rep-like genes are displayed in orange, virion structural genes shown in green, other identifiable viral genes shown in pink, other genes in grey. GenBank accession numbers are displayed above the virus name.

Notably, a large CRESS genome (CRESS virus isolate ctdh33, associated with rhabditid nematodes that were serially cultured from a soil sample) encoded three separate genes with structural homology (HHpred probability scores 97-99%) to STNV capsid (Fig 4C). The three predicted STNV capsid homologs in the nematode virus are highly divergent from one another, with only 28-30% amino acid similarity, but also highly divergent from other amino acid sequences in GenBank. A possible explanation for this observation is that the capsid gene array is the result of gene duplication events.

To further characterize the soil nematode virus, the three candidate capsid proteins were overexpressed in combination in human-derived 293TT cells (Buck et al., 2004). Particles apparently composed of all three capsid proteins were able to sediment in Optiprep gradients. Further, analysis by negative stain electron microscopy (EM) revealed ~50 nm particles (Fig S5A). Interestingly these particles were found to contain nonspecific nuclease-resistant RNA (Fig S5B). This supports the concept that CRESS virus capsid proteins might theoretically be re-targeted to packing RNA virus genomes or vice versa.

CRESS genomes ctba10, ctcc19, ctbj26, ctcd34, and ctbd1037 (ranging from 3.5 - 6.2 kb in length) also each encode two divergent capsid gene homologs (Fig. 4D, Fig S6). Single genomes encoding multiple capsid genes with related but distinct amino acid sequences have been observed in RNA viruses (Agranovsky et al., 1995) and giant dsDNA viruses (Schulz et al., 2017), but we believe that this is the first time it has been reported in ssDNA viruses.

Two related large CRESS viruses (ctdb796 and ctce741) encode capsid proteins similar to those of bacilladnaviruses (Fig 4G,I). Interestingly, the Rep genes of the two viruses do not show close similarity to known bacilladnavirus Reps and are instead similar to the Reps of certain unclassified CRESS viruses, suggesting that CRESS ctdb796 and CRESS ctce741 are representatives of a new hybrid CRESS virus family.

Two other CRESS virus genomes (isolates ctca5 and ctgh4) encode capsid genes that show amino acid similarity to distinct groups of icosahedral T=3 ssRNA virus capsids (Makino et al., 2013) (tombus- and tombus-like viruses), but not to cruciviruses or bacilladnaviruses (Fig 4A,F, Fig. 5). Further, a 6.6 kb CRESS virus (isolate ctbd466) (Fig 4H) was found to encode a gene with some similarity to the capsid region of the polyprotein of two newly described ssRNA viruses (ciliovirus and brinovirus (Fig. S7B) (Makino et al., 2013, Greninger and DeRisi, 2015). Protein fold predictor Phyre^2^ (Kelley et al., 2015) showed a top hit (58% confidence) for the capsid protein of a norovirus (ssRNA virus with T=3 icosahedral capsid) for isolate ctbd466 (see GenBank: AXH73946).

**Figure 5:**
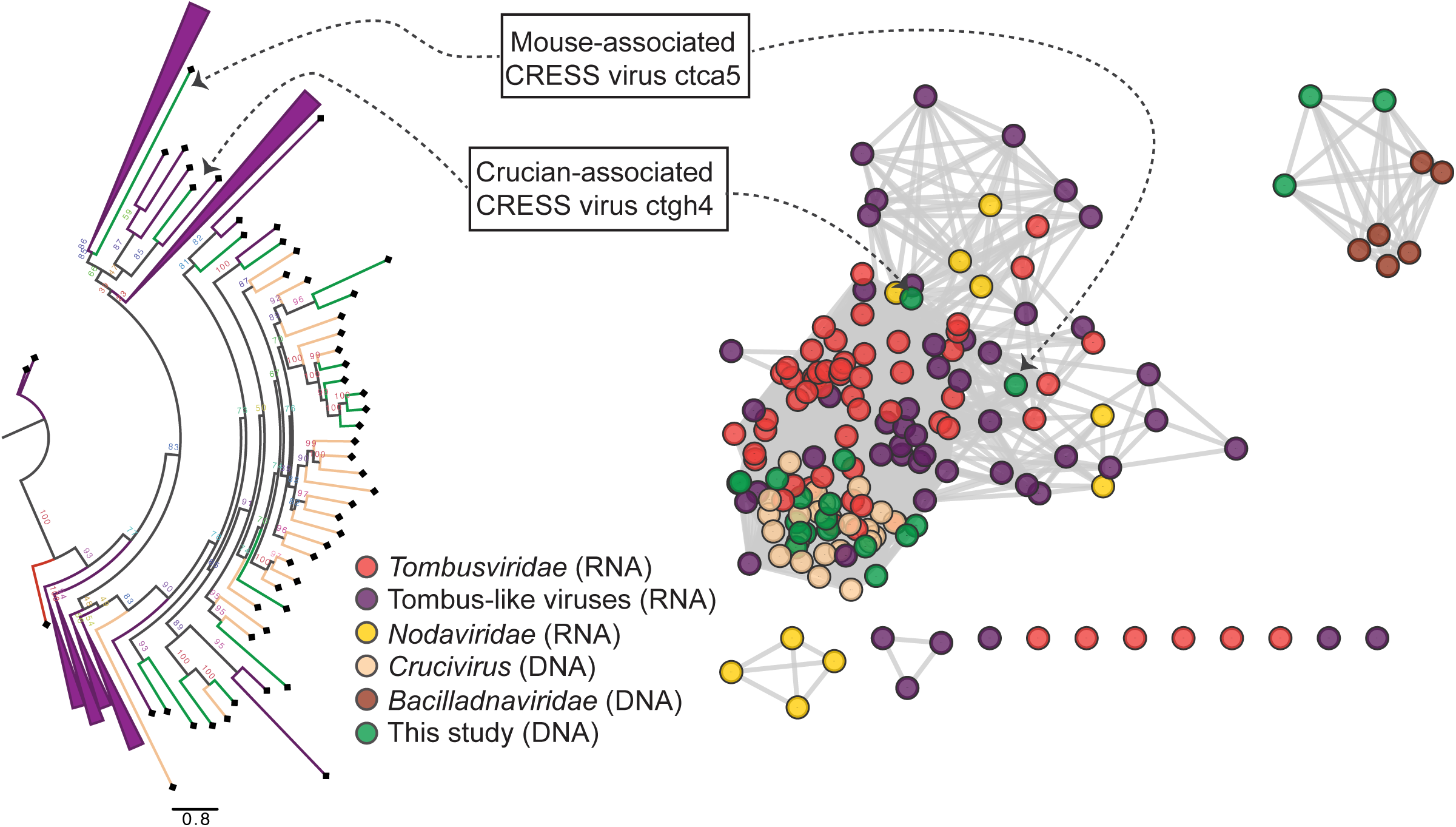
RNA virus capsid-like proteins. Sequence similarity network generated with EFI-EST (E value cutoff of 10-5) showing capsid protein sequences of select ssRNA viruses (*Nodaviridae*, *Tombusviridae*, tombus-like viruses) and ssDNA viruses (*Bacilladnaviridae* and crucivirus) together with protein sequences from DNA virus genomes observed in the present study with predicted structural similarity to an RNA virus capsid protein domain (PDB: 2IZW). Predicted capsid proteins for CRESS virus ctca5 and CRESS virus ctgh4 have no detectible similarity to any known DNA virus sequences. On the left, a phylogenetic tree representing the large cluster is displayed. Collapsed branches consist of *Tombusviridae*, tombus-like viruses, and *Nodaviridae* capsid genes.

Two CRESS genomes (ctbe30 and ctbc27) from separate Rhesus macaque stool samples combine Rep genes specific to CRESS viruses with several genes specific to inoviruses, including inovirus-like capsid genes, which encode proteins that form a filamentous virion (Fig 4B, J). The bacteriophage families *Inoviridae* and *Microviridae* are ssDNA viruses that replicate via the rolling circle mechanism, but they are not considered conventional CRESS viruses because they infect prokaryotes and do not encode Rep genes with CRESS-like sequences. Other inovirus-like genes encoded in the ctbe30 and ctbc27 genomes include homologs of zonular occludens toxin (ZOT, a packaging ATPase) and RstB (a DNA-binding protein required for host genome integration) (Falero et al., 2009) (Fig 4B, J). TBLASTX searches using ctbe30 and ctbc27 sequences yielded large segments of similarity to various bacterial chromosomes (e.g., GenBank accession numbers AP012044 and AP018536), presumably representing integrated prophages. This suggests that ctbb30 and ctbc27 represent a previously undescribed bacteria-tropic branch of the CRESS virus supergroup.

Viral genomes discussed in this section were validated by aligning individual reads back to the contigs followed by visual inspection. No disjunctions were detected, indicating that illegitimate recombinations are not evident (see Fig. S7C for an example).

### Network analysis of genetic “dark matter” demonstrates conservation of gene sequence and genome structure

We defined potential viral “dark matter” in the survey as circular contigs with no hits with E values <1×10^−5^ in BLASTX searches of a database of viral and plasmid proteins. We posited that leveraging sequence similarity networks would be useful both for analyzing groups of gene homologs and for discerning which gene combinations tended to be present on related circular genomes. To categorize the 609 dark matter elements based on their predicted proteins, we used pairwise comparison with EFI-EST. A majority of translated gene sequences could be categorized into dark matter protein clusters (DMPCs) containing four or more members (Fig. 6A). Further, groups of related dark matter elements (i.e. dark matter genome groups (DMGGs)), much like viral families, could be delineated by the presence of a conserved, group-specific marker gene. For example, DMPC1 can be thought of as the marker gene for DMGG1. Certain DMPCs tend to co-occur on the same DMGG. For instance, DMPC7 and DMPC17 ORFs are always observed in genomes with a DMPC1 ORF (i.e., DMGG1) (Fig. 6B). This *pro tempore* categorization method is useful for visualizing the data, but we stress that is not necessarily taxonomically definitive.

**Figure 6:**
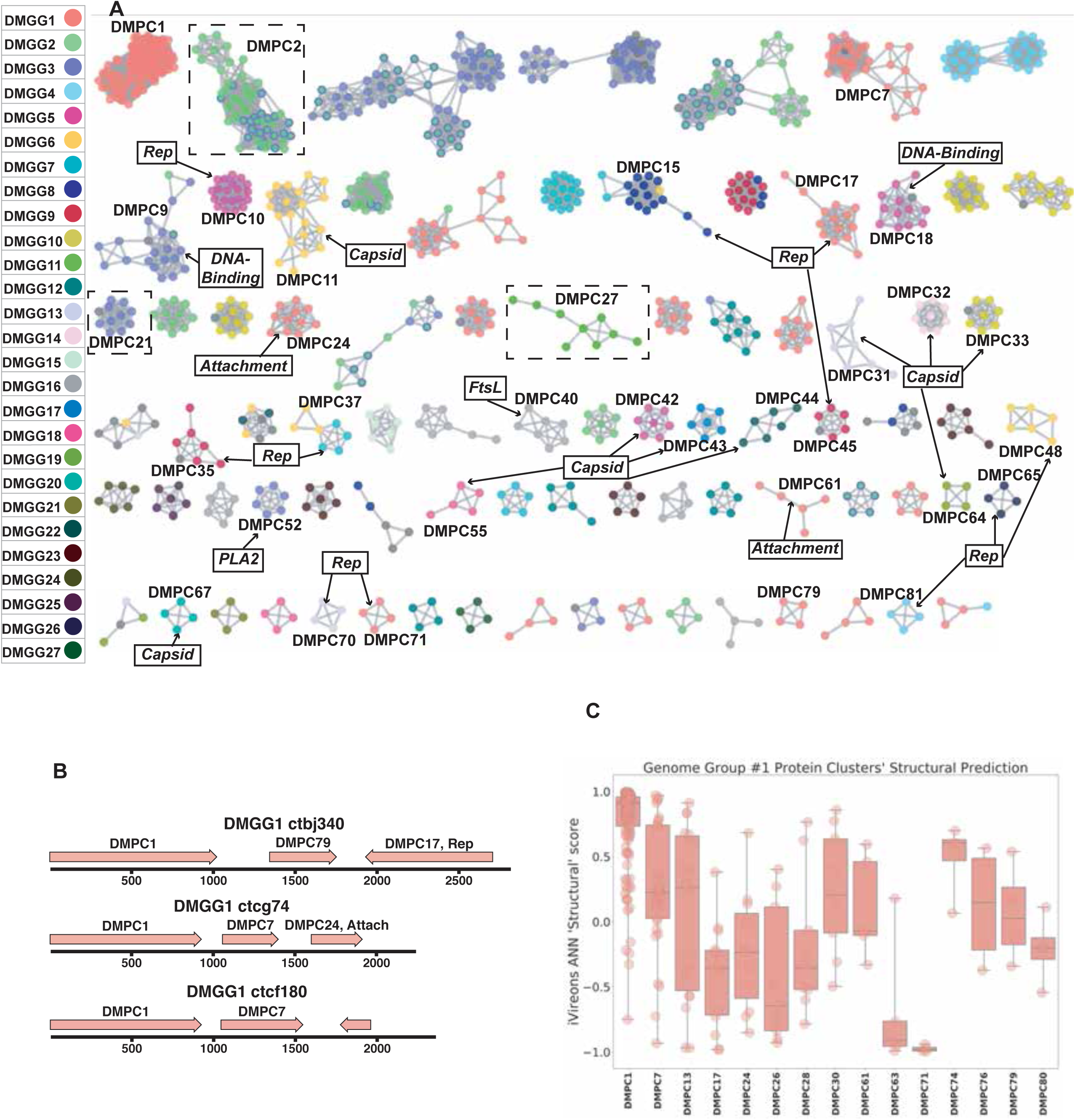
Dark matter analysis. (A) Sequence similarity network analysis for genes from dark matter circular sequences (minimum cluster size = 4). Clusters are colored based on assigned dark matter genome group (DMGG). Structural predictions from HHpred are indicated (>85% probability). *Rep* = rolling circle replicases typical of CRESS viruses or ssDNA plasmids. *Capsid* = single-jellyroll capsid protein. *Attachment* = cell attachment proteins typical of inoviruses. *DNA-Binding* = DNA-binding domain. *PLA2* = phospholipase A2. *FtsL* = FtsL-like cell division protein. Clusters that contain a representative protein that was successfully expressed as a virus-like particle are outlined by a dashed rectangle (See Fig. 7). (B) Maps of three examples of DMGG1 with DMPCs labeled (linearized for display). (C) DMGG1 iVireons ‘structure’ score summary by protein cluster. Scores range from −1 (unlikely to be a virion structural protein) to 1 (likely to be a virion structural protein). Additional iVireons score summaries can be found in Fig. S8.

HHpred, was again employed to make structural predictions for these data (Zimmermann et al., 2018). Instead of querying individual sequences, alignments were prepared using MAFFT (Katoh and Standley, 2013) for each major DMPC to identify conserved residues and increase sensitivity. Then, each alignment was used for an HHpred query. The results indicate that ten DMPCs are likely viral capsid proteins and 11 are rolling circle replicases (Fig. 6A).

While most of the circular dark matter in the survey could be characterized using these methods, dark matter contigs represent a small remaining fraction in some samples (Fig. S9).

### Cell culture expression of candidate “dark matter” capsids yields particles

In contrast to viral genes such as Rep with conserved enzymatic functions, sequences of the capsid genes are often poorly conserved, even within a given viral family (Buck et al., 2016). Moreover, it appears that capsid proteins have arisen repeatedly through capture and modification of different host cell proteins (Krupovic and Koonin, 2017). This makes it challenging to detect highly divergent capsid proteins using alignment-based approaches or even structural modelling. We therefore turned to an alignment-independent approach known as iVireons, an artificial neural network trained by comparing alignment-independent variables between a large set of known viral structural proteins and known non-structural proteins (Seguritan et al., 2012) (https://vdm.sdsu.edu/ivireons/). As an example of the approach, iVireons scores for DMPCs associated with DMGG1 are shown in Fig. 6C. Other sets of iVireons scores can be seen in Fig. S8.

Of the 17 DMGGs for which HHPRED did not identify capsid genes, iVireons predicted that ten contain at least one DMPC predicted to encode some type of virion structural protein (median score of cluster >0.70). This allowed us to generate the testable hypothesis that some of these predicted structural proteins would form virus-like particles (VLPs) if expressed in cell culture.

A subset of predicted capsid proteins were expressed in 293TT cells and/or in *E. coli* and subjected to size exclusion chromatography. Electron microscopic analysis showed that several of the predicted capsid proteins formed roughly spherical particles, whereas a negative control protein did not form particles (Fig. 7). Although the particles were highly irregular, the DMGC11 isolate ctgh70 preparation was found to contain nuclease-resistant nucleic acids, consistent with nonspecific encapsidation. The results suggest that, in multiple cases, we were able to experimentally confirm that iVireons correctly predicted the identity of viral capsid proteins.

**Figure 7:** Expression of putative capsid proteins. Images taken by negative stain electron microscopy. Genome maps are linearized for display purposes. Expressed genes are colored green. iVireons scores are listed in parentheses. (A-C) Images represent virus-like particles from iVireons-predicted viral structural genes. (D) Merkel cell polyomavirus small T antigen (a viral non-structural protein) is shown as a negative control.

## Discussion

Massive parallel DNA sequencing surveys characterizing microbial communities typically yield a significant fraction of reads that cannot be mapped to known genes. The present study sought to provide the research community with an expanded catalog of viruses with circular DNA genomes associated with humans and animals, as well as a means to characterize future datasets. We hope that the availability of this expanded viral sequence catalog will facilitate future investigation into associations between viral communities and disease states. Our annotation pipeline, Cenote-Taker, can be accessed via http://www.cyverse.org/discovery-environment. The CyVerse version of Cenote-Taker can readily annotate circular or linear DNA viruses. RNA viruses with polyproteins or frameshifts will require *post hoc* manual editing.

At the present time, GenBank’s RefSeq database includes complete sequences for 8112 viral genomes, most of which fit into 131 families recognized by the International Committee on Taxonomy of Viruses (ICTV) (King et al., 2018). Similarly, the IMG/VR database contains over 14,000 circular virus genomes from hundreds of studies, though some of these appear to be redundant with each other and are not comprehensively annotated (Paez-Espino et al., 2019). The current study, which focused on circular DNA viruses with detergent-resistant capsids, found 2,514 new complete circular genomes. The availability of these comprehensively annotated genomes in GenBank contributes new information and understanding to a broad range of established, emerging, and previously unknown taxa. Figure 3 shows dozens of potential family-level groupings within the unclassified CRESS virus supergroup. Sequences from this study contribute to 40 of such groupings and constitute the only members of seven groups. There are also 192 singleton CRESS sequences that could establish many additional family-level groups.

Although small ssDNA viruses are ubiquitous, they are often overlooked in studies that only characterize sequences that are closely related to reference genomes. In addition, ssDNA is not detected by some current DNA sequencing technologies unless second-strand synthesis (such as the RCA approach used in the current study) is conducted.

While many of the viruses discovered in this study appear to be derived from prokaryotic commensals, it is important to note that bacteriophages can contribute to human and animal diseases by transducing toxins, antimicrobial resistance proteins, or genes that alter the physiology of their bacterial hosts (Waldor and Mekalanos, 1996). Furthermore, interaction between animal immune systems and bacteriophages appears to be extensive (Hodyra-Stefaniak et al., 2015).

Over 100 distinct human anellovirus sequences were found in human blood. Anelloviruses have yet to be causally associated with any human disease, but this study indicates that we are likely still just scratching the surface of the sequence diversity of human anelloviruses. It will be important to fully catalog this family of viruses to address the field’s general assumption that they are harmless.

Several of the CRESS viruses detected in this study are larger than any other CRESS virus genomes that have been described previously. In some cases, the larger size of these genomes may have been enabled by a process involving capsid gene duplication events. Further, CRESS virus acquisition of T=3 capsids from ssRNA *Nodaviridae* and *Tombusviridae* families has been previously suggested as the origin of bacilladnaviruses (Kazlauskas et al., 2017) and cruciviruses (Steel et al., 2016, Dayaram et al., 2016, Roux et al., 2013, Krupovic et al., 2015), respectively. We present evidence of additional independent recombination events between CRESS viruses and ssRNA viruses and ssDNA bacteriophages. In light of these findings, it should be reiterated that only DNA (not RNA) was sequenced in our approach, so DNA/RNA *in silico* false recombination does not seem plausible. These data suggest that CRESS viruses are at the center of a tangled evolutionary history of viruses in which genomes change not just via gradual point mutations but also through larger scale recombination and hybridization events.

It is likely that some dark matter sequences detected in this study share a common ancestor with known viruses but are too divergent to retain discernable sequence similarity. In some cases, the dark matter circles may represent a more divergent segment of a virus with a multipartite genome. Alternatively, some of these sequences likely represent entirely new viral lineages that have not previously been recognized.

## Methods

### LEAD CONTACT AND MATERIALS AVAILABILITY

Further information and requests for resources and reagents should be directed to and will be fulfilled by the Lead Contact, Chris Buck

### EXPERIMENTAL MODEL AND SUBJECT DETAILS

Human-derived 293TT cells (female) were generated in-house (Buck et al., 2004). The cells were not authenticated for this study.

T7 Express lysY/I^q^ *E. coli* were purchased from New England Biolabs.

### METHOD DETAILS

#### Sample collection and sequencing

De-identified human swabs and tissue specimens were collected under the approval of various Institutional Review Boards (Supplemental Table 1). Animal tissue samples were collected under the guidance of various Animal Care and Use Committees.

Nematodes were cultured out of soil samples collected in Bethesda, Maryland, USA on OP50-Seeded NGM-lite plates (*C. elegans* kit, Carolina Biological Supply).

Viral particles were concentrated by subjecting nuclease-digested detergent-treated lysate to ultracentrifugation over an Optiprep step gradient, as previously described https://ccrod.cancer.gov/confluence/display/LCOTF/Virome (Peretti et al., 2015). Specifically, for each sample, no more than 0.5 g of solid tissue was minced finely with a razorblade. Alternatively, no more than 500 µl of liquid sample was vortexed for several seconds. Samples were transferred to 1.5ml siliconized tubes. The samples were resuspended in 500 µl Dulbecco’s PBS and Triton X-100 (Sigma) detergent was added to a final concentration of 1% w/v. 1 µl of Benzonase (Sigma) was added. Samples were vortexed for several seconds. Samples were incubated in a 37°C water bath for 30 minutes, with brief homogenising using a vortex every 10 minutes. After incubation, NaCl was added to the samples to a final concentration of 0.85M. Tubes were spun for 5 minutes at 5000g. Resulting supernatants were transferred to a clean siliconized tube. Supernatant-containing tubes were spun for an additional 5 minutes at 5000g. Resulting supernatants were added to iodixanol/Optiprep (Sigma) step gradients in ultracentrifuge tubes (Beckman: 326819) (equal volumes 27%, 33%, 39% iodixanol with 0.8M NaCl; total tube volume, including sample, ~5.1ml). Ultracentrifuge tubes were spun at 55,000rpm for 3.5 hours (Beckman: Optima L-90K Ultracentrifuge). After spin, tubes were suspended over 1.5ml siliconized collection tubes and pierced at the bottom with 25G needle. Six fractions of equal volume were collected drop-wise from each ultracentrifuge tube.

From each fraction, 200 µl was pipette to a clean siliconized tube for virus particle lysis and DNA precipitation. To disrupt virus particles, 50 µl of a 5X master mix of Tris pH 8 (Invitrogen, final conc. 50mM), EDTA (Invitrogen, final conc. 25mM), SDS (Invitrogen, final conc. 0.5%), Proteinase K (Invitrogen, final conc. 0.5%), DTT (Invitrogen, final conc. 10mM) was added and mixed by pipetting up and down. Samples were heated at 50°C for 15 minutes. Then, proteinase K was inactivated for 10 minutes at 72°C. To the 250 µl of sample, 125 µl of 7.5M ammonium acetate was added and mixed by vortexing. Then, 975 µl of 95% ethanol was added and mixed by pipetting. This was incubated at room temperature for 1 hour. Then, the samples were transferred to a 4°C fridge overnight.

Samples were then restored to ambient temperature. Then, samples were spun for 1 hour at 20,000g in a temperature-controlled tabletop centrifuge set to 21°C. Supernatant was aspirated, and 500 µl ethanol was added to each pellet. Pellets were resuspended by flicking. Then, samples were spun for 30 minutes at 20,000g in a temperature-controlled tabletop centrifuge set to 21°C. Supernatant was aspirated, and samples were spun once more at 20,000g for 3 minutes. Remaining liquid was carefully removed with a 10 µl micropipette. Tubes were left open and air dried for at least 10 minutes. DNA from individually collected fractions of the gradient was amplified by RCA using phi29 polymerase (TempliPhi, Sigma) per manufacturer’s instructions. While we expected most viral particles to travel to the middle of the gradient based on previous experiments, RCA was conducted on individual fractions spanning the gradient, in an attempt to detect viruses with different biophysical properties (Kauffman et al., 2018). Pooled, amplified fractions were prepared for Illumina sequencing with Nextera XT kits. Then libraries were sequenced with Illumina technology on either MiSeq or NextSeq500 sequencers. Contigs were assembled using SPAdes with the ‘plasmid’ setting. Circularity was confirmed by assessing assembly graphs using Bandage (Wick et al., 2015).

#### Analysis of brain samples

Brain samples were initially analyzed by Optiprep gradient purification, RCA amplification, and deep sequencing, as described above. JC polyomavirus, which has previously been reported in brain samples (Chalkias et al., 2018), can display high buoyancy in Optiprep gradients (Geoghegan et al., 2017). Fractions from near the top of the Optiprep gradient were subjected to an alternative method of virion enrichment using microcentrifuge columns (Pierce) packed with 2 ml of Sepharose 4B Bead suspension (Sigma) exchanged into PBS. Fractions were clarified at 5000 x g for 1 minute, and 200 µl of clarified extract was loaded onto the gel bed. The column was spun at 735 x g and the eluate was digested with proteinase K, ethanol-precipitated, and subjected to RCA. No additional viral sequences were detected by this method.

The brain samples were also subjected to confirmatory analysis by RNA sequencing. RNA was extracted from brain tissues with Qiagen Lipid Tissue RNeasy Mini Kit and subjected to human ribosomal RNA depletion with Thermo RiboMinus. The library was prepared with NEBNext Ultra™ II Directional RNA Library Prep Kit for Illumina and subjected to massive parallel sequencing on the Illumina HiSeq platform (see BioProject PRJNA513058).

#### Cenote-Taker, Virus Discovery and Annotation Pipeline

Cenote-Taker, a bioinformatics pipeline written for this project and fully publicly available on CyVerse, was used for collection and detailed annotation of each circular sequence. The flow of the program can be described as follows:

1. Identifies and collects contigs (assembled with SPAdes) larger than 1000 nts
2. Predicts which contigs are circular based on overlapping ends
3. Determines whether circular contig has any ORFs of 80 AA or larger or else discards sequence
4. Uses BLASTN against GenBank ‘nt’ database to disregard any circular sequences that are >90% identical to known sequences across a >500 bp window
5. Uses Circlator (Hunt et al., 2015) to rotate circular contigs so that a non-intragenic start codon of one of the ORFs will be the wrap point
6. Uses BLASTX against a custom virus + plasmid database (derived from GenBank ‘nr’ and RefSeq) to attempt to assign the circular sequence to a known family
7. Translates each ORF of 80 AA or larger
8. Uses RPS-BLAST to predict function of each ORF by aligning to known NCBI Conserved Domains
9. Generates a tbl file of RPS-BLAST results
10. Takes ORFs without RPS-BLAST hits and queries the GenBank ‘nr viral’ database with BLASTP
11. Generates a tbl file of BLASTP results
12. Takes ORFs without any BLASTP hits and queries HHblits (databases: uniprot20, pdb70, scop70, pfam_31, NCBI_CD)
13. Generates a tbl file of HHblits results
14. Complies with a GenBank request to remove annotations for ORFs-within-ORFs that do not contain conserved sequences
15. Combines all tbl files into a master tbl file
16. Generates a unique name for each virus based on taxonomic results
17. Generates properly formatted fsa and tbl files in a separate directory
18. Uses tbl2asn to make gbf (for viewing genome maps) and sqn files (for submission to GenBank)

The source code can be found at: https://github.com/mtisza1/Cenote-Taker

This work utilized the computational resources of the NIH HPC Biowulf cluster. (http://hpc.nih.gov).

Genome maps were drawn, and multiple sequence alignments were computed and visualized using MacVector 16.

#### Anelloviruses

Analysis of linear contigs in the survey found many instances of recognizable viral sequences. One noteworthy example were anelloviruses, where many contigs terminated near the GC-rich stem-loop structure that is thought to serve as the origin of replication. This segment of the anellovirus genome is presumably incompatible with the short read deep sequencing technologies used in this study. Nearly complete anellovirus genomes, defined as having a complete ORF1 gene and at least 10-fold depth of coverage, were also deposited in GenBank (Table S1).

#### GenBank Sequences

Amino Acid sequences from ssDNA viruses were downloaded in June 2018 based on categories in the NCBI taxonomy browser. As many sequences in GenBank are from identical/closely related isolates, all sequences were clustered at 95% AA ID using CD-HIT (Fu et al., 2012).

#### Sequence Similarity Networks

Amino acid sequences from GenBank (see above) and this study were used as queries for HHsearch (the command-line iteration of HHpred) against PDB, PFam, and CDD. Sequences that had hits in these databases of 80% probability or greater were kept for further analyses. Note that capsid protein models for some known CRESS virus families have little, if any, similarity to other capsid sequences and have not been determined (e.g. *Genomoviridae* and *Smacoviridae*) and were therefore not displayed in networks. Models used: (CRESS virus capsids network:5MJF_V, 3R0R_A, 5MJF_Ba, 4V4M_R, 4BCU_A, PF04162.11, 5J37_A, 5J09_C, 3JCI_A, cd00259, PF04660.11, PF03898.12, PF02443.14, pfam00844); (CRESS virus Rep network:4PP4_A, 4ZO0_A, 1M55_A, 1UUT_A, 1U0J_A, 1S9H_A, 4R94_A, 4KW3_B, 2HWT_A, 1L2M_A, 2HW0_A, PF08724.9, PF17530.1, PF00799.19, PF02407.15, pfam08283, PF12475.7, PF08283.10, PF01057.16, pfam00799); (*Microviridae*/*Inoviridae* replication-associated protein: 4CIJ_B, 4CIJ_C, PF05155.14, PF01446.16, PF11726.7, PF02486.18, PF05144.13, PF05840.12); (*Microviridae* capsid: 1M06_F, 1KVP_A, PF02305.16); (*Anelloviridae* ORF1: PF02956.13); (*Inoviridae* ZOT: 2R2A_A, PF05707.11).

#### Phylogenetic Trees

Sequences from this study and GenBank were grouped by structural prediction using HHpred. Then, sequences were compared by EFI-EST to generate clusters with a cut-off of 1×10^−5^. Sequences from these clusters were then extracted and aligned with PROMALS3D (Pei and Grishin, 2014) using structure guidance, when possible. Structures used: (*Microviridae* MCP: 1KVP); (CRESS virus capsid STNV-like: 4V4M); (CRESS virus capsid circo-like: 3JCI); (*Inoviridae* ZOT: 2R2A); (CRESS virus Rep: 2HW0) (CRESS virus/RNA virus S Domain capsid: 2IZW). The resulting alignments were used to build trees with IQ-Tree with automatic determination of the substitution model and 1000 ultrafast bootstraps (Nguyen et al., 2015). Models used: (*Microviridae* MCP: Blosum62+F+G4); (*Microviridae* Rep I: Blosum62+I+G4); (*Microviridae* Rep II: LG+I+G4); (*Microviridae* Rep III: VT+I+G4); (CRESS virus/RNA virus S Domain capsid: Blosum62+F+G4); (*Circoviridae* capsid: VT+F+G4); (CRESS virus capsid STNV-like: VT+F+G4); (*Inoviridae* ZOT: VT+I+G4); (*Anelloviridae* ORF1: VT+F+G4). Trees were visualized with FigTree (http://tree.bio.ed.ac.uk/software/figtree/) and iTOL (Letunic and Bork, 2019).

#### Expressing Potential Viral Structural Proteins in human 293TT cells

Codon-modified expression constructs encoding the three predicted STNV-like capsid proteins from the nematode virus (CRESS virus isolate ctdh33) were designed according to a previously reported “as different as possible” algorithm https://ccrod.cancer.gov/confluence/display/LCOTF/Pseudovirus+Production. 293TT cells were transfected with combinations of STNV capsid-like protein expression constructs for roughly 48 hours. Cells were lysed in a small volume of PBS with 0.5% Triton X-100 or Brij-58 and Benzonase (Sigma). After several hours of maturation at neutral pH, the lysate was clarified at 5000 x g for 10 min. The clarified lysate was loaded onto a 27-33-39% Optiprep gradient in PBS with 0.8 M NaCl. Gradient fractions were collected by bottom puncture of the tube and screened by PicoGreen nucleic acid stain (Invitrogen), BCA, and SDS-PAGE analysis. Electron microscopic analysis was then performed. Expression in 293TT cells of some “dark matter” virus capsids was attempted but not successful in any case.

#### Expressing Potential Viral Structural Proteins in *E. coli*

Several genes that were identified by iVireons as being potential viral structural proteins were cloned into plasmids with a T7 polymerase-responsive promoter. Plasmids were transfected into T7 Express lysY/I^q^ *E. coli*, which express T7 polymerase under the induction of IPTG. Bacteria were grown at 37°C in LB broth until OD600 = 0.5. Flasks were cooled to room temperature, IPTG was added to 1 mM, and cultures were shaken at room temperature for approximately 16 hours. Cells were then pelleted for immediate processing.

Total protein was extracted with a BPER (Pierce) and nuclease solution. Then, virion-sized particles were enriched from the clarified lysate using size exclusion chromatography with 2% agarose beads https://ccrod.cancer.gov/confluence/display/LCOTF/GelFiltration. Fractions were analyzed using Coomassie-stained SDS-PAGE gels for presence of a unique band corresponding to the expressed protein. Fractions of interest were analyzed using negative stain electron microscopy.

#### Electron Microscopy

Five µl samples were adsorbed onto a carbon-deposited copper grid for one minute. Sample was then washed 5 times on water droplets then stained with 0.5% uranyl acetate for 1 second. The negatively stained samples were examined on a FEI Tecnai T12 transmission electron microscope.

#### Sequencing

Illumina sequencing was conducted at the CCR Genomics Core at the National Cancer Institute, NIH, Bethesda, MD 20892.

### DATA AND CODE AVAILABILITY

All reads and annotated genomes associated with this manuscript can be found on NCBI BioProject Accessions PRJNA393166 and PRJNA396064.

Cenote-Taker, the viral genome annotation pipeline, can be used by interested parties on the Cyverse infrastructure: http://www.cyverse.org/discovery-environment.

### ADDITIONAL RESOURCES

Relevant protocols on lab website: https://ccrod.cancer.gov/confluence/display/LCOTF/Virome

## Acknowledgements

This research was supported [in part] by the Intramural Research Program of the NIH, National Cancer Institute.

We would like to acknowledge the GenBank team at NCBI for productive discussion about their viral genome submission requirements and facilitation of annotated genome deposition.

## Author Contributions

M.J.T. conceived, performed, and validated experiments, developed methodologies and software, performed analysis, curated data, wrote and revised the manuscript, and visualized data. D.V.P., N.L.W., B.S., A.P., and Y.S.P conceived and performed experiments and revised the manuscript. G.J.S. and S.R.K. developed software and revised the manuscript. A.M.S. performed analysis, conceived of experiments, and revised the manuscript. A.V. performed analysis, visualized data, and revised the manuscript. P.A.P., D.H.M., P.M.M., J.L.W., B.M., J.M.B., S.P.R., B.R., J.D., B.A.T., O.P., J.T., and S.M.R. provided resources and reviewed the manuscript. B.B. and M.B.S. developed methods and software and reviewed the manuscript. C.B.B. conceived of experiments, developed methods and software, supervised and administered the project, and revised the manuscript.

---

**Supplemental Figure 1:**
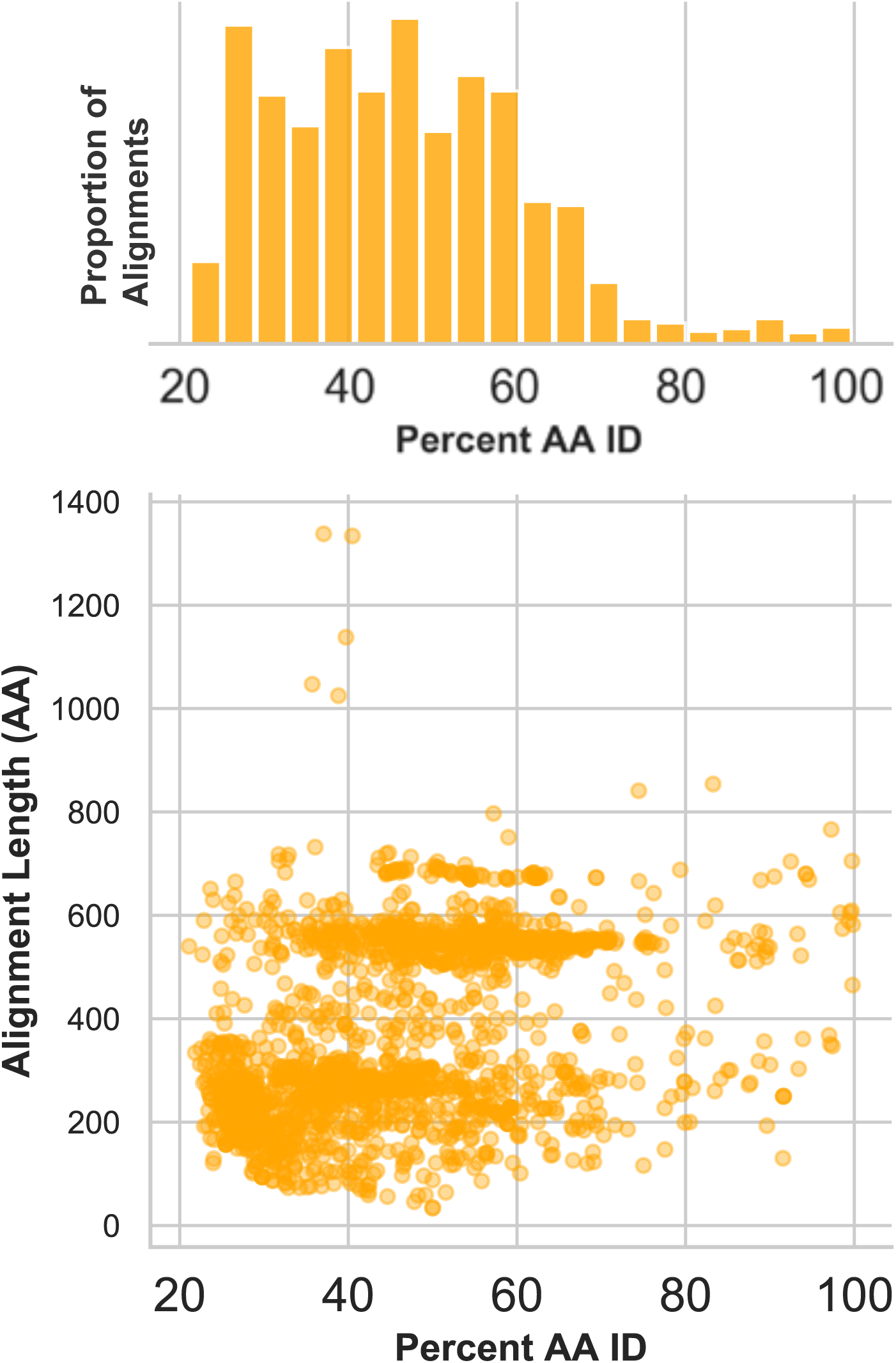
Divergence of proteins encoded by circular contigs. BLASTX summary of each circular DNA molecule recovered from virus enriched samples. Sequences were queried against a database of viral and plasmid sequences. Only hits with E values < 10^−5^ were plotted. Here, BLASTX only reports the most significant stretch of amino acid sequence from each circular contig, and, therefore, other regions of each contig can be assumed to be equally or less conserved.

**Supplemental Figure 2:**
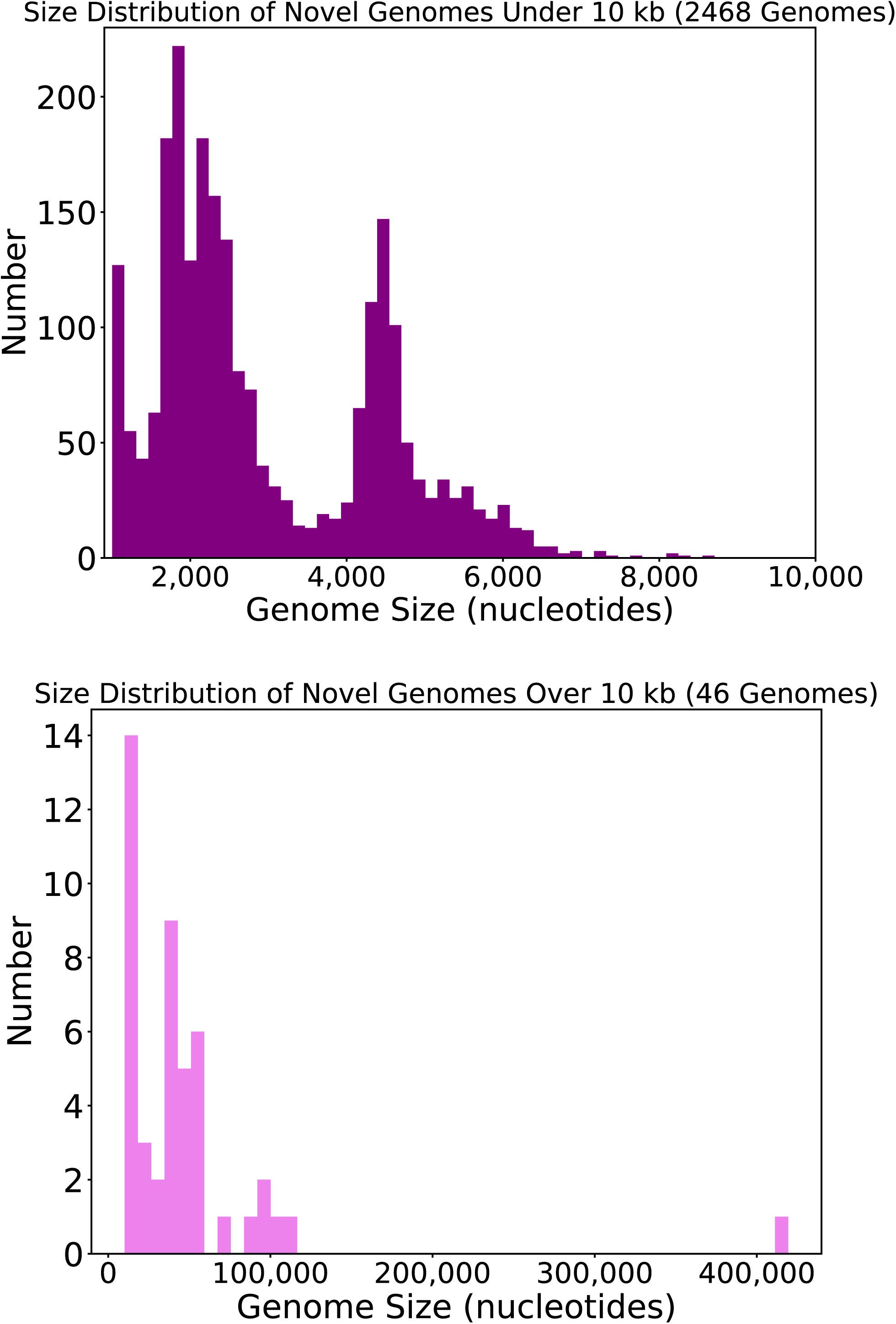
Size distribution of circular DNA sequences from this study. Length, in nucleotides, of circular DNA sequences representing putative viral genomes from this study.

**Supplemental Figure 3:**
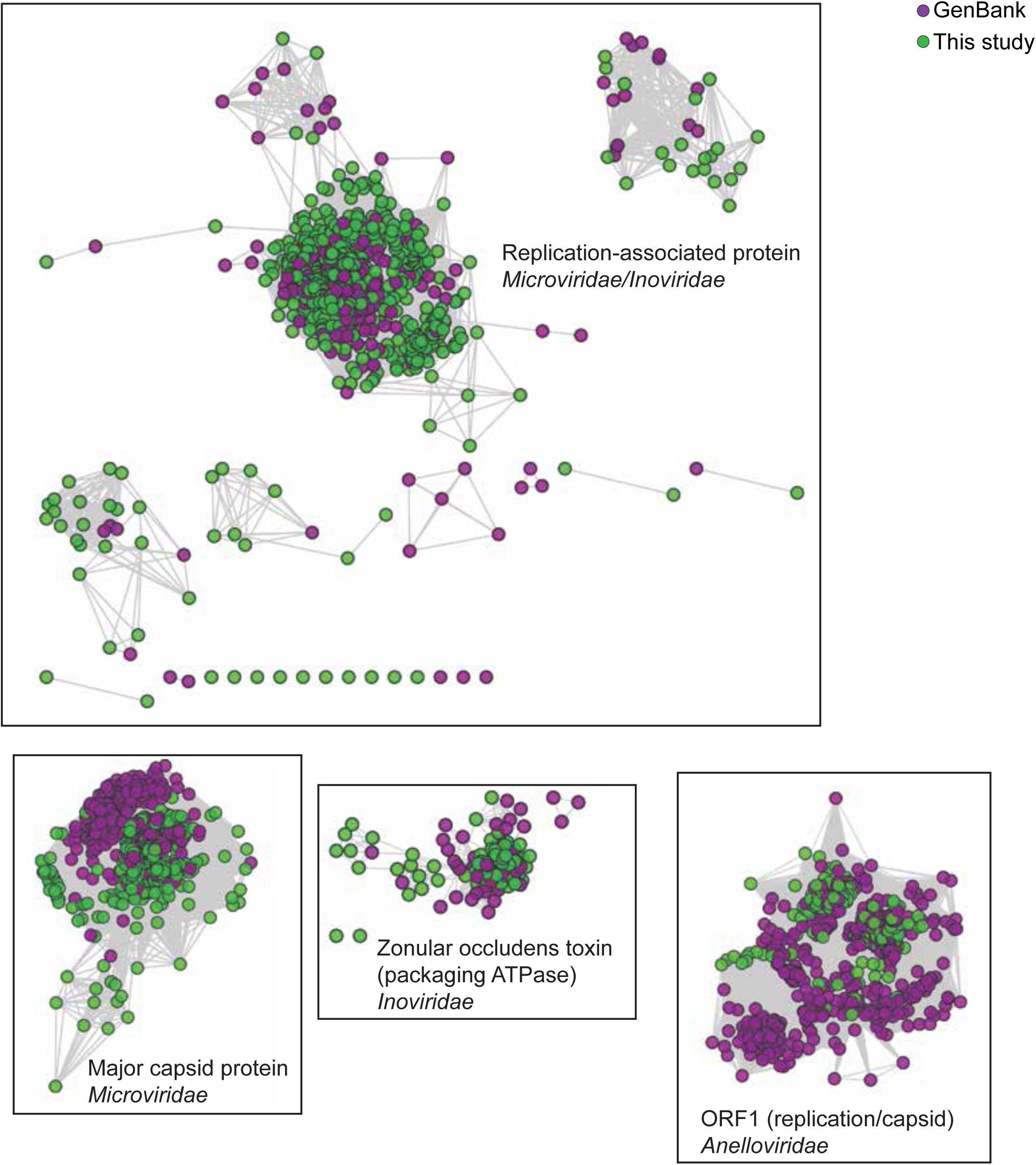
Network Analysis of additional viral hallmark genes. Depiction of additional viral hallmark genes from this study and GenBank as sequence similarity networks. E value cutoff = 10^−5^. See Figure 2 and Methods.

**Supplemental Figure 4:**
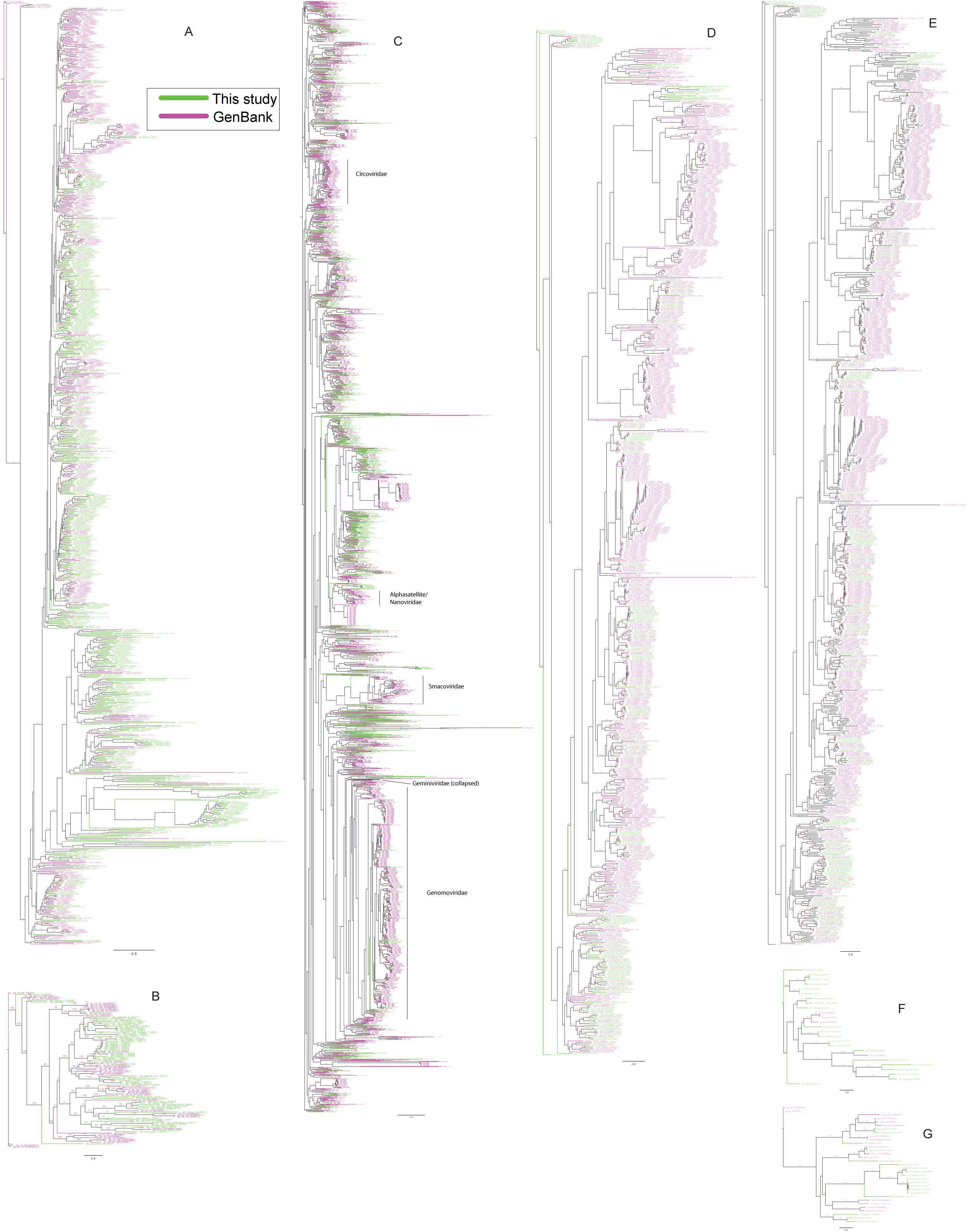
Phylogenetic trees of viral hallmark genes. Sequences were aligned with PROMALS3D using structure guidance when possible. Trees were drawn using IQ-Tree with automatic determination of substitution model. See methods. Branches are labeled with bootstrap percent support after 1000 ultrafast bootstrapping events. (A) *Microviridae* major capsid protein. (B) *Inoviridae* zonular occludens toxin. (C) CRESS virus Rep. (D) *Anelloviridae* ORF1 (E) *Microviridae*/*Inoviridae* Replication-associated protein I. (F) *Microviridae*/*Inoviridae* Replication-associated protein II. (G) *Microviridae*/*Inoviridae* Replication-associated protein III.

**Supplemental Figure 5:**
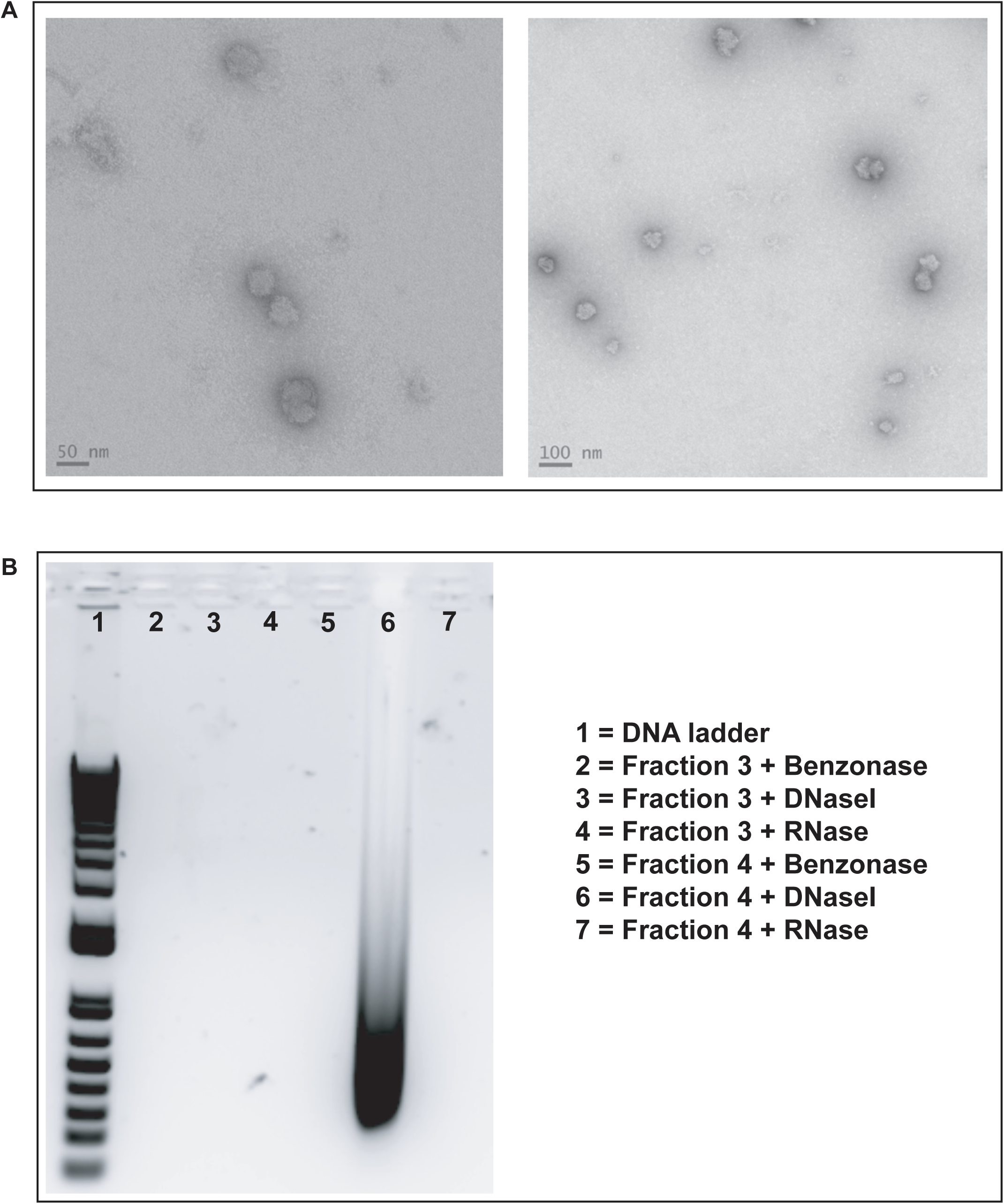
Nematode-associated virus capsid expression. Putative capsid proteins from CRESS virus ctdh33 associated with rhabditid nematodes (see Fig. 3) were expressed in a human cell line. (A) Negative Stain EM images of virus-like particles resulting from the expression of all three genes predicted to encode STNV-like capsid proteins. (B) Agarose gel electrophoretic analysis of nucleic acids extracted from ctdh33 VLPs shows that purified VLPs from fraction 4 of the Optiprep gradient contain encapsidated RNA. Note that Benzonase digests both DNA and RNA, DnaseI only digests DNA and Rnase only digests RNA.

**Supplemental Figure 6:**
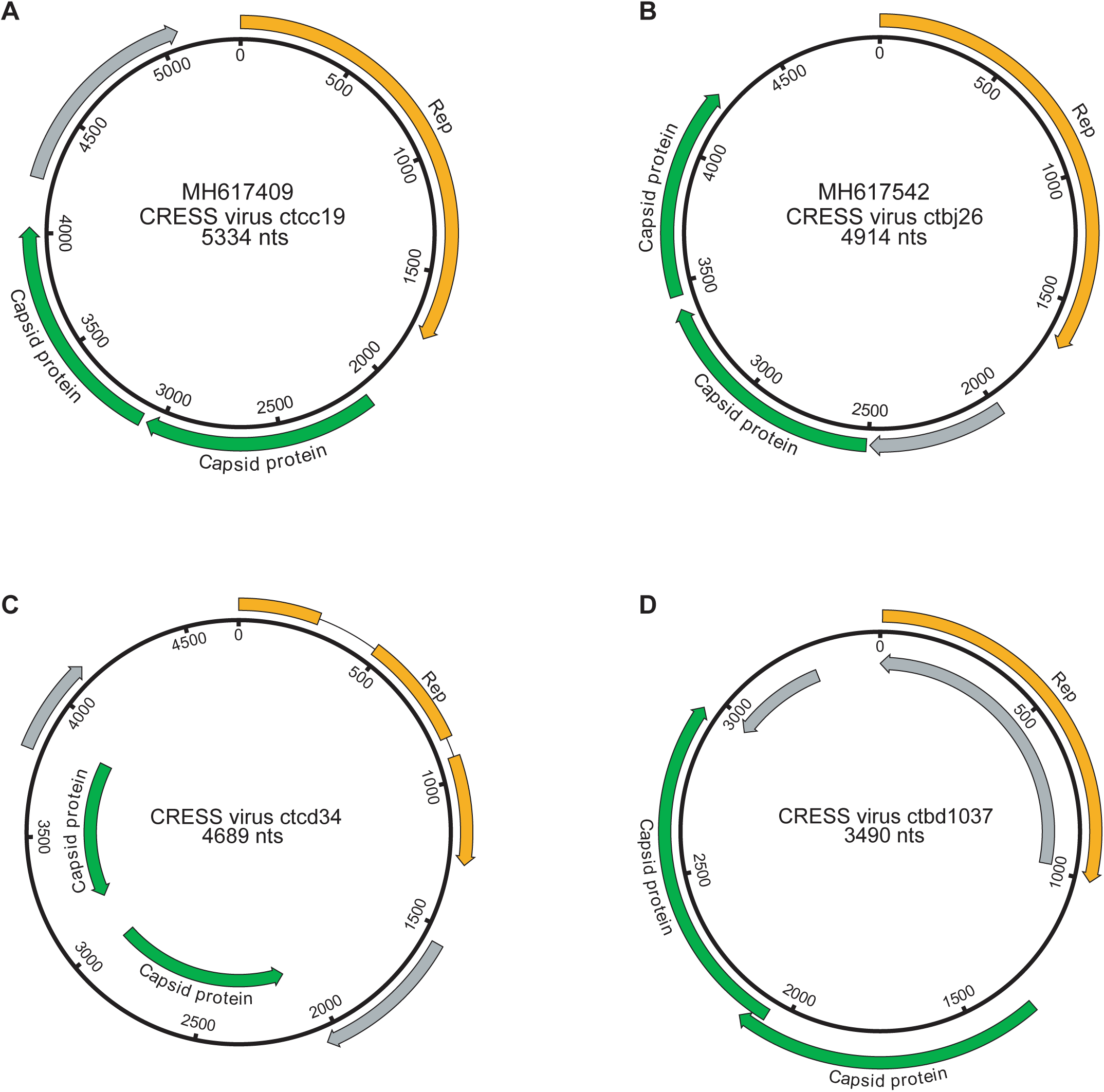
Genome maps of additional CRESS genomes encoding multiple capsid genes. CRESS Rep genes are shown in orange, predicted virion structural genes in green, other candidate genes in gray.

**Supplemental Figure 7:**
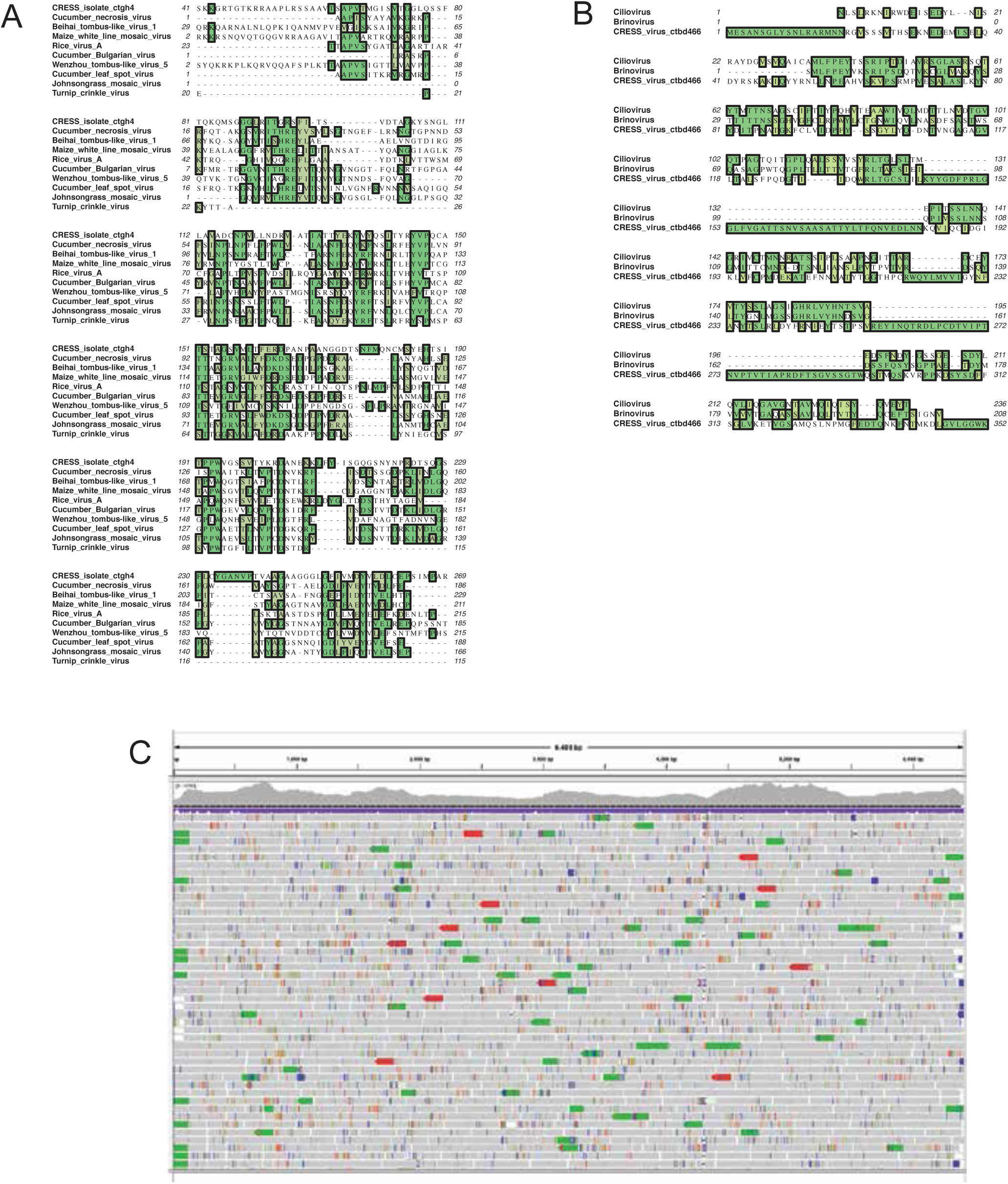
Validation of proteins with predicted similarity to RNA virus capsid proteins. (A) First order neighbors for Crucian-associated CRESS virus ctgh4 capsid protein were extracted from the network shown in Figure 5 and aligned using Muscle. (B) The same approach was applied to CRESS virus ctbd466 capsid protein. (C) A visualization (Integrative Genomics Viewer) of a read alignment to CRESS virus isolate ctca5. The visualization shows no evidence of artifactual chimerization in the contig assembly process.

**Supplementary Figure 8 (related to Figure 6):**
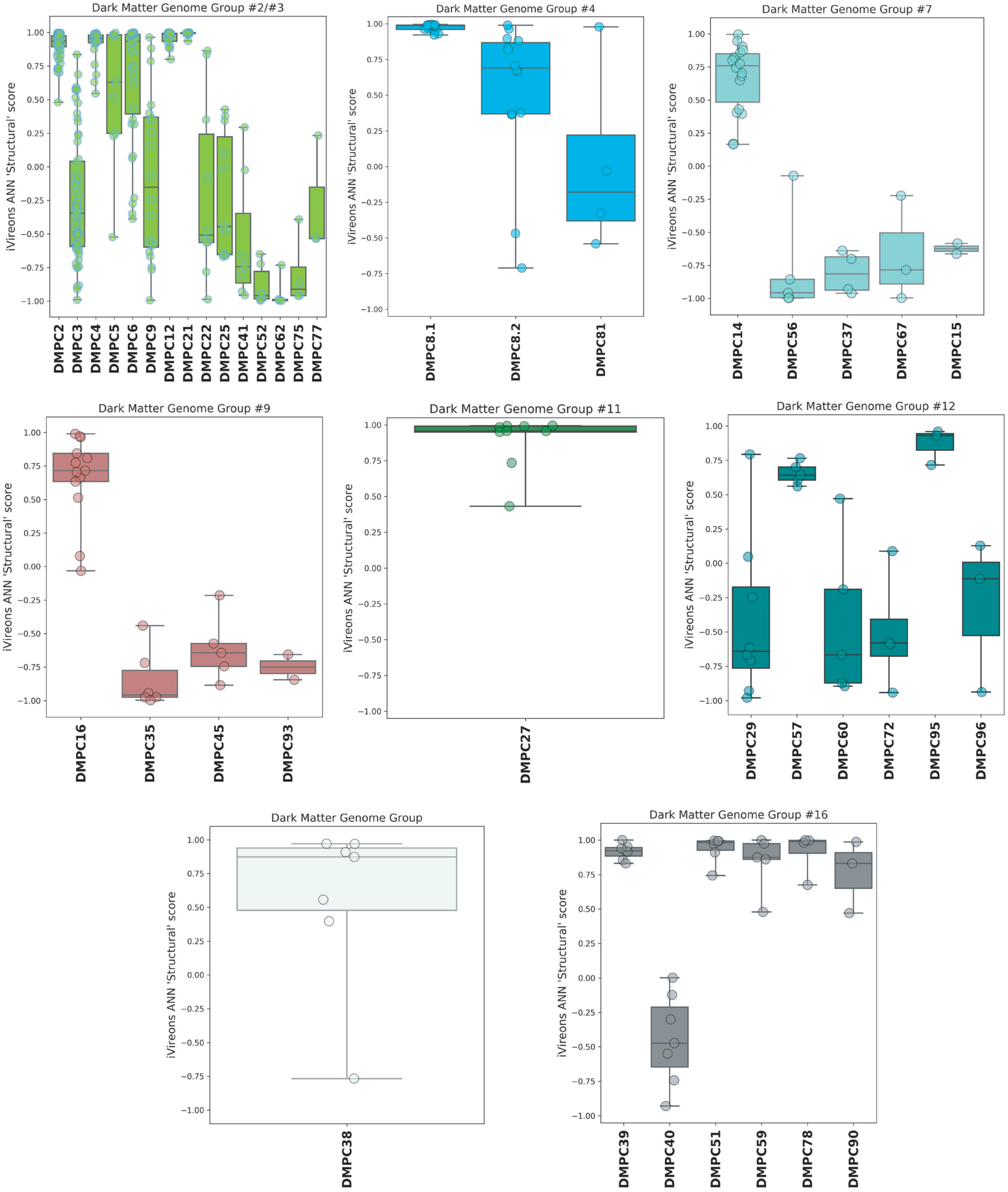
iVireons scores of DMGGs with candidate viral structural gene(s) Box-and-whisker plots of iVireons ‘Structural’ scores for individual DMPCs (numbers on x-axes) grouped by DMGG. Scores (y-axes) range from −1 (unlikely to be a virion structural protein) to 1 (likely to be a virion structural protein). DMGG2 and DMGG3 have been combined due to inferred chimerism.

**Supplemental Figure 9:**
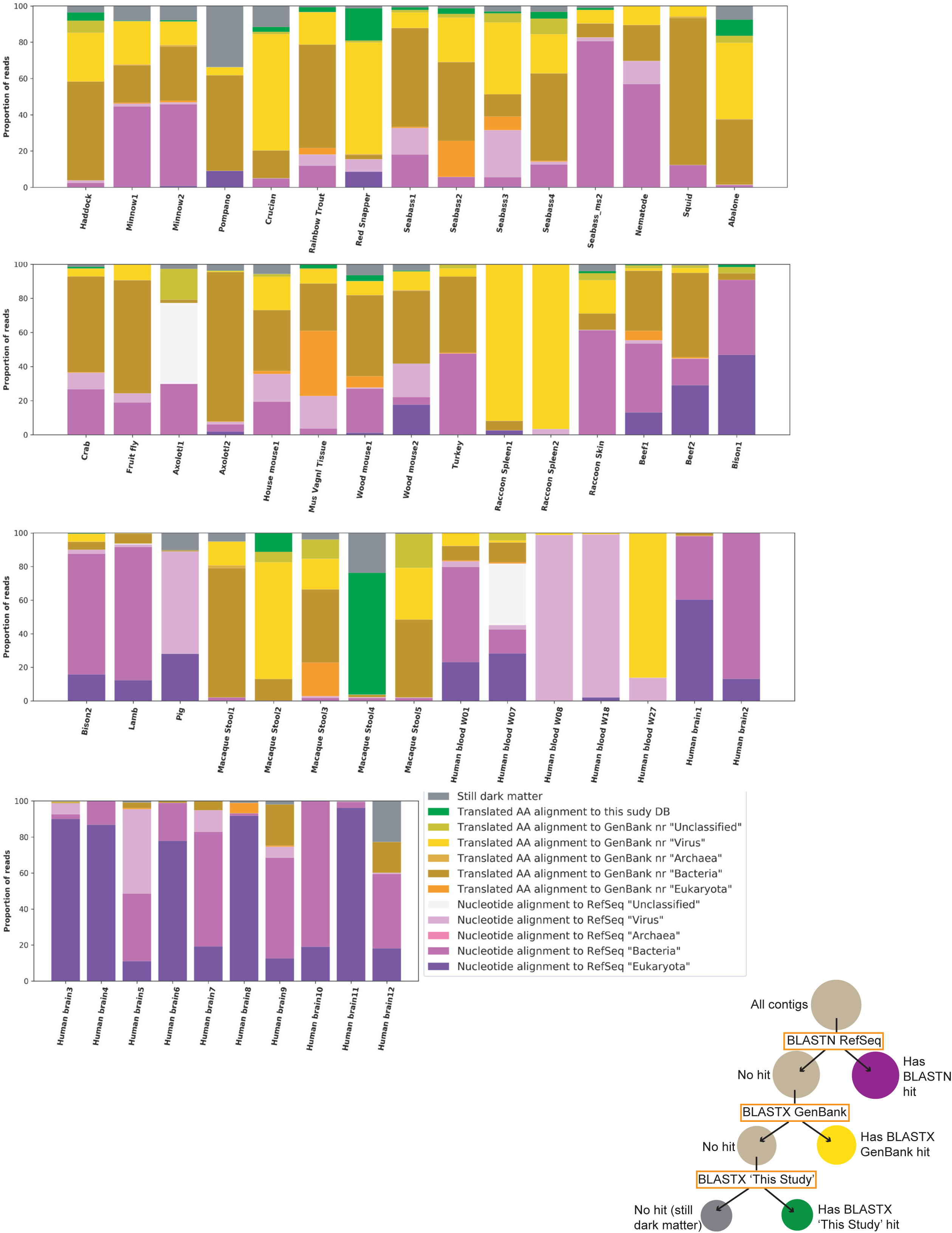
Sample characterization by iterative BLAST Searches. Contigs of over 1000 nts from each sample were subject to iterative BLAST searches. First, BLASTN was performed against the RefSeq database. Contigs without hits were then queried by BLASTX against all of GenBank ‘nr’ database. Contigs without hits were then queried by BLASTX against a database of proteins from genomes reported in this study. The proportion of total reads mapping to each contig was calculated and used for this plot. Individual inspection of contigs shows that most hits in the ‘Translated AA alignment to GenBank ‘nr’ “Bacteria”’ were likely plasmid or prophage proteins. The proportions of hits in each category are sensitive to stringency settings and to which databases are chosen for the analysis. The key aims of the figure are to display the proportion of reads the current survey rendered classifiable and the fraction of remaining dark matter reads in various samples.

**Table S1.**
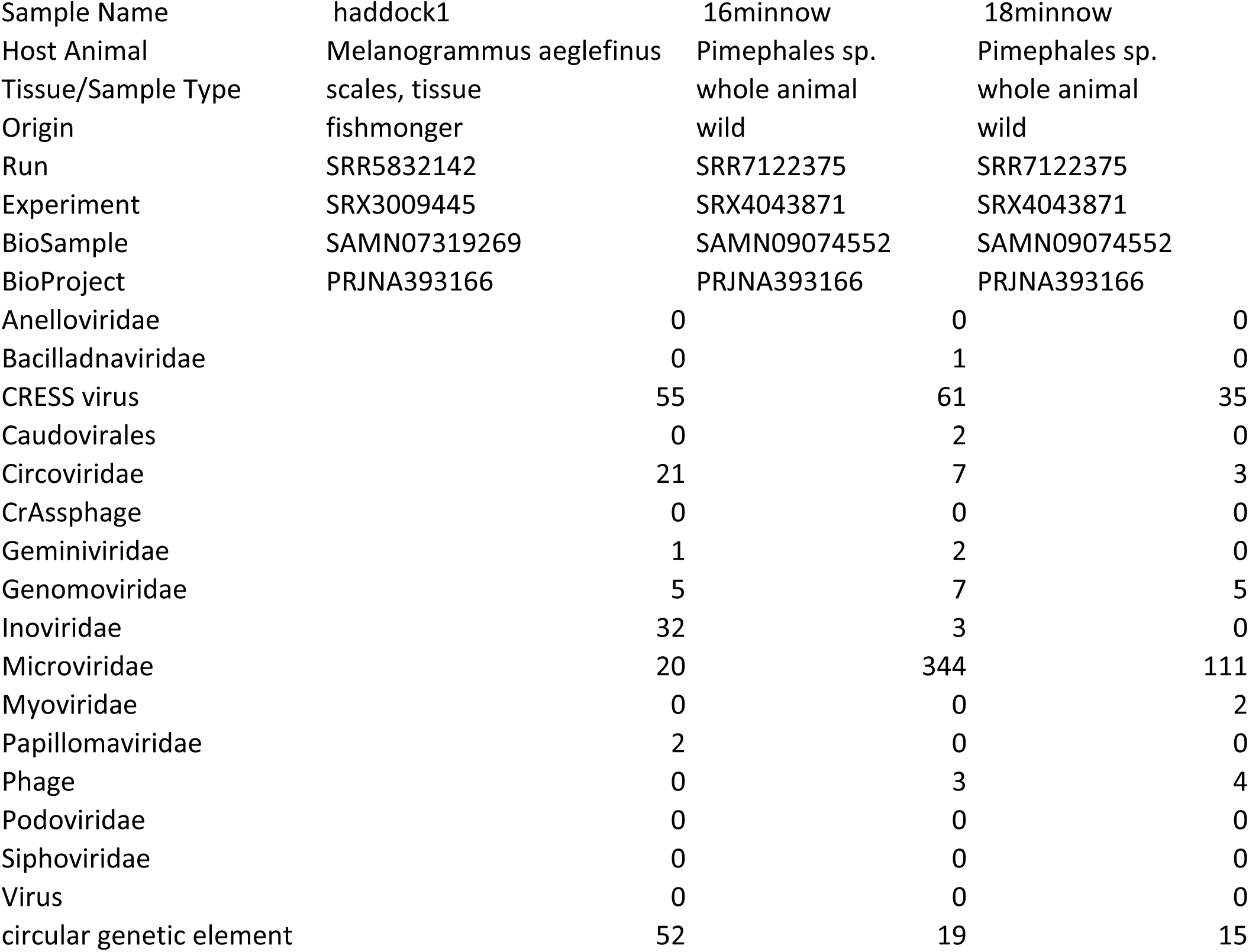
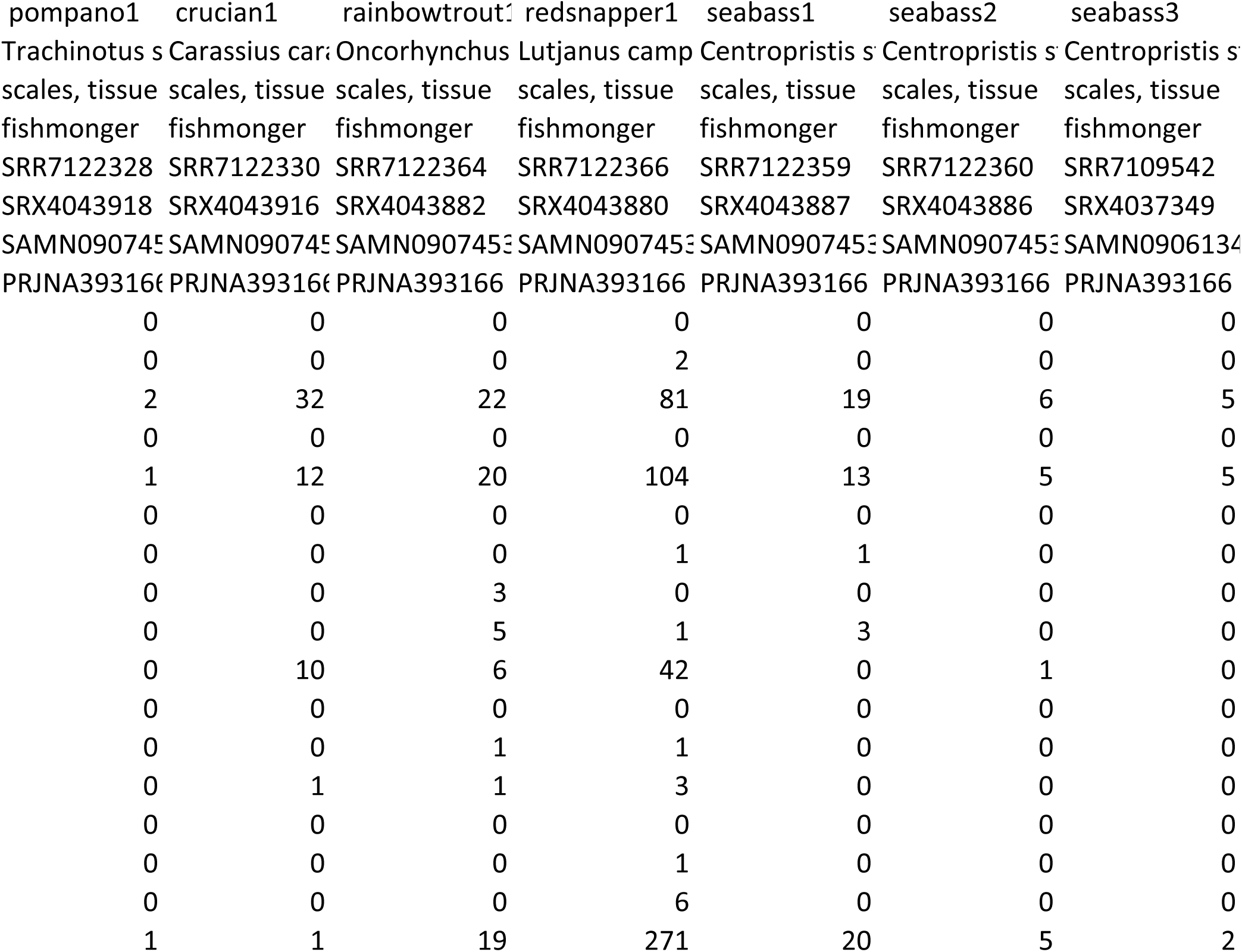
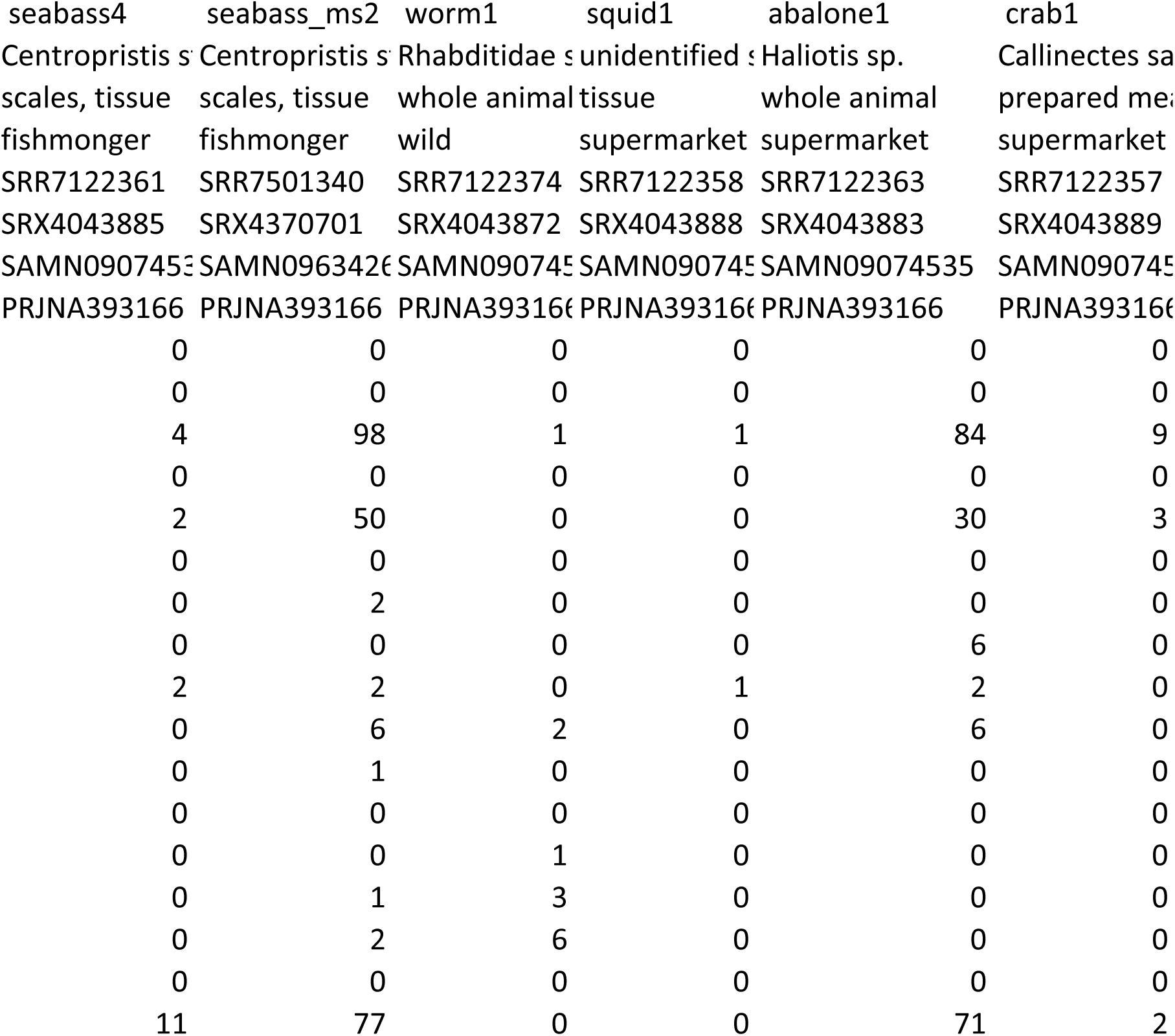
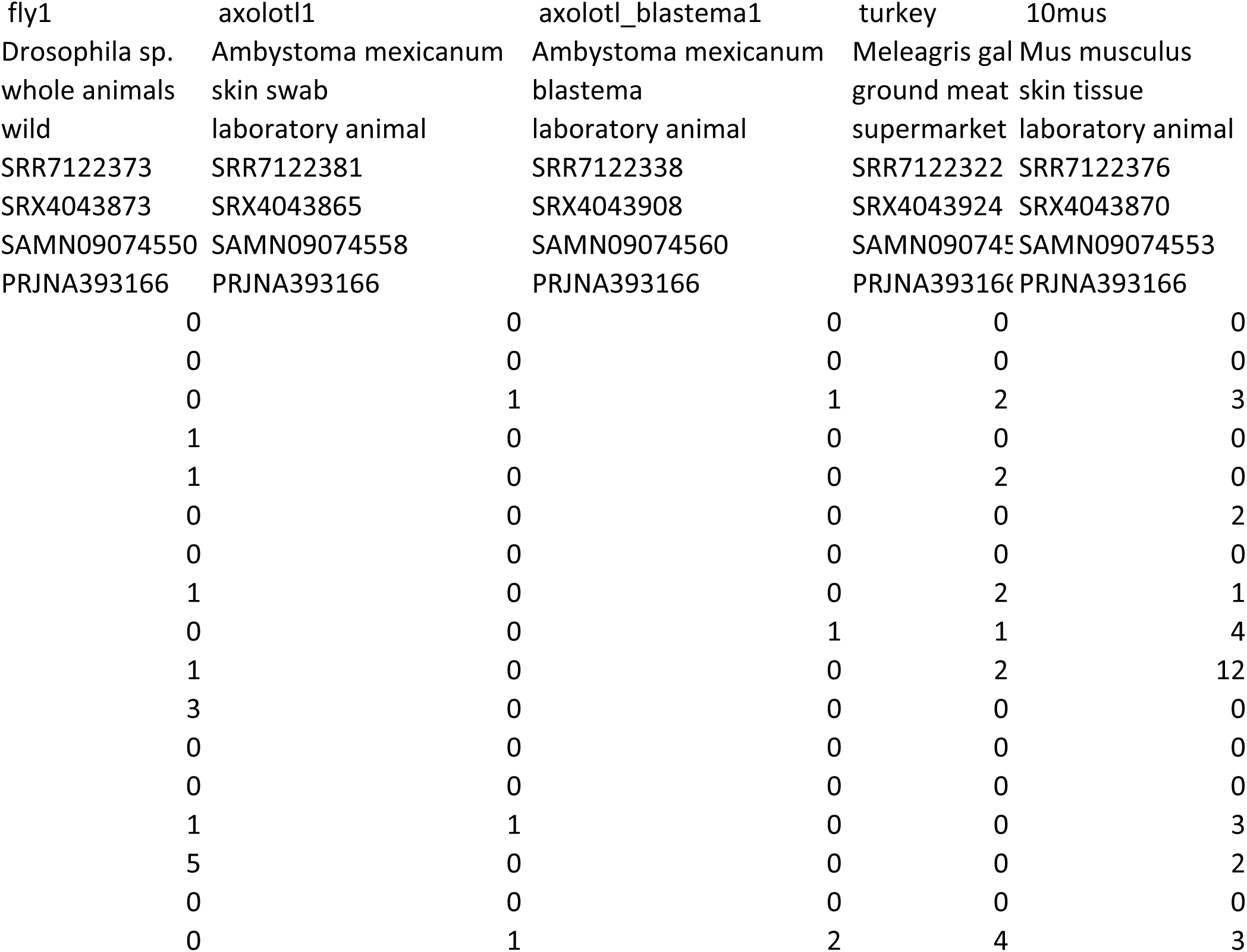
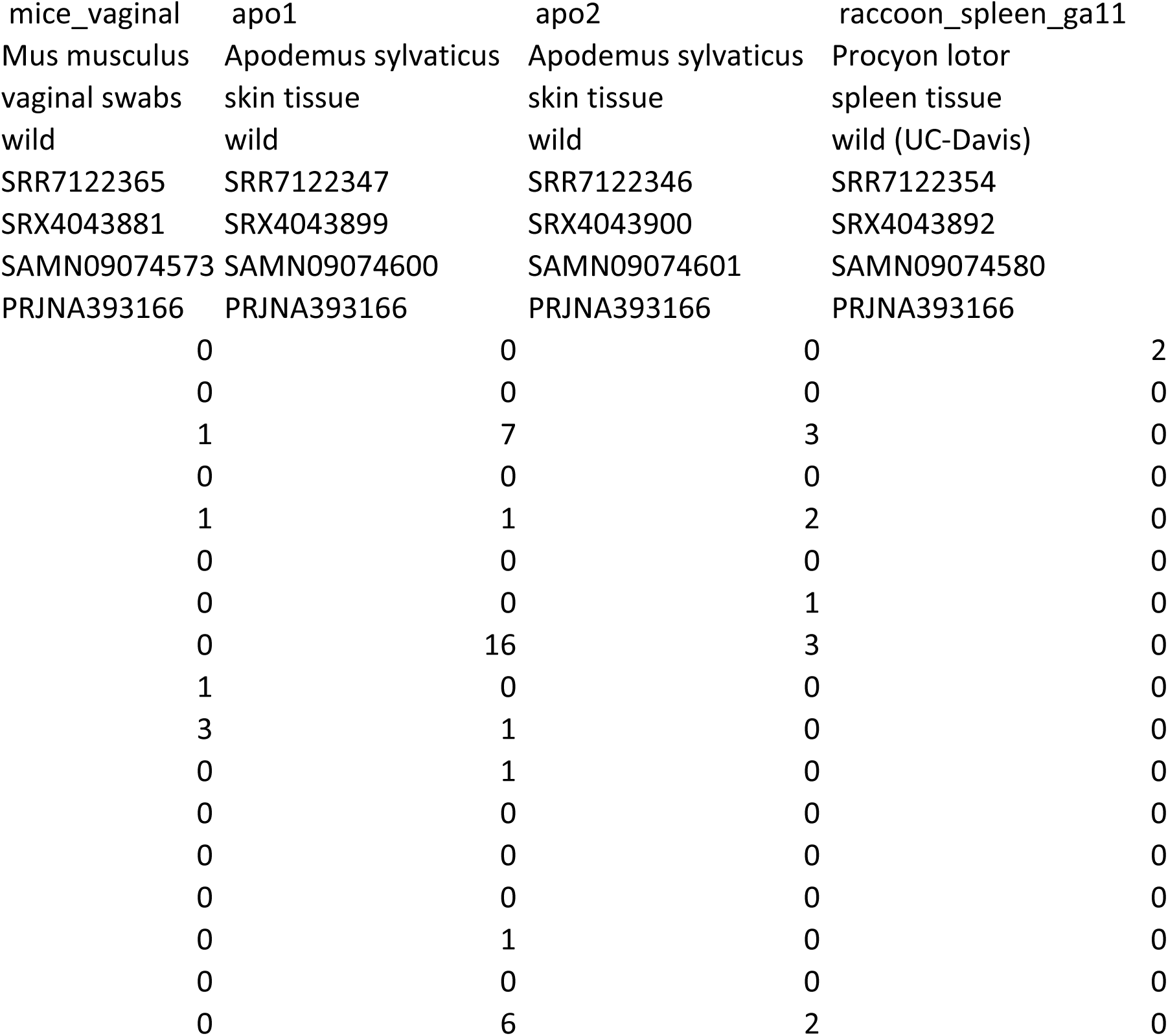
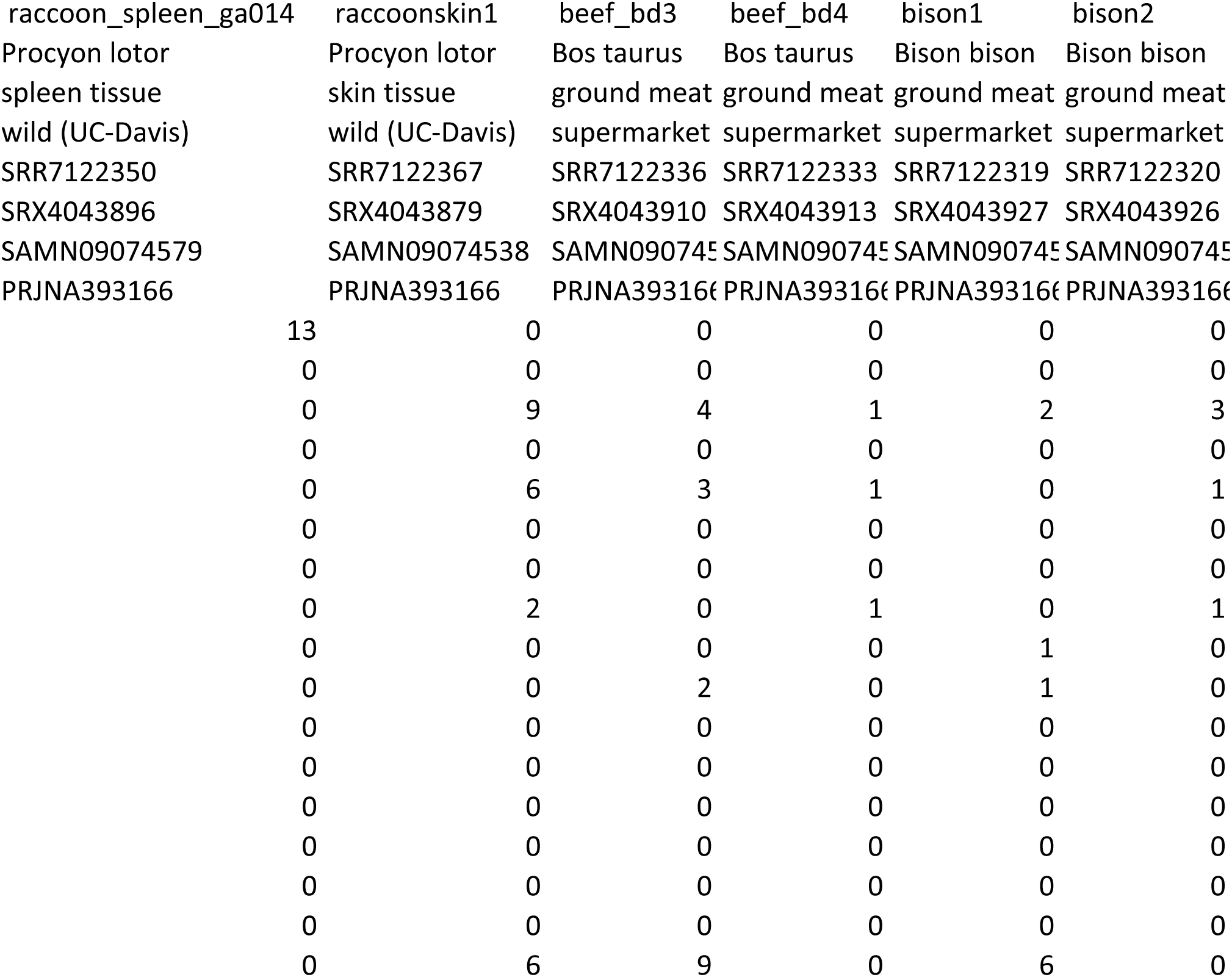
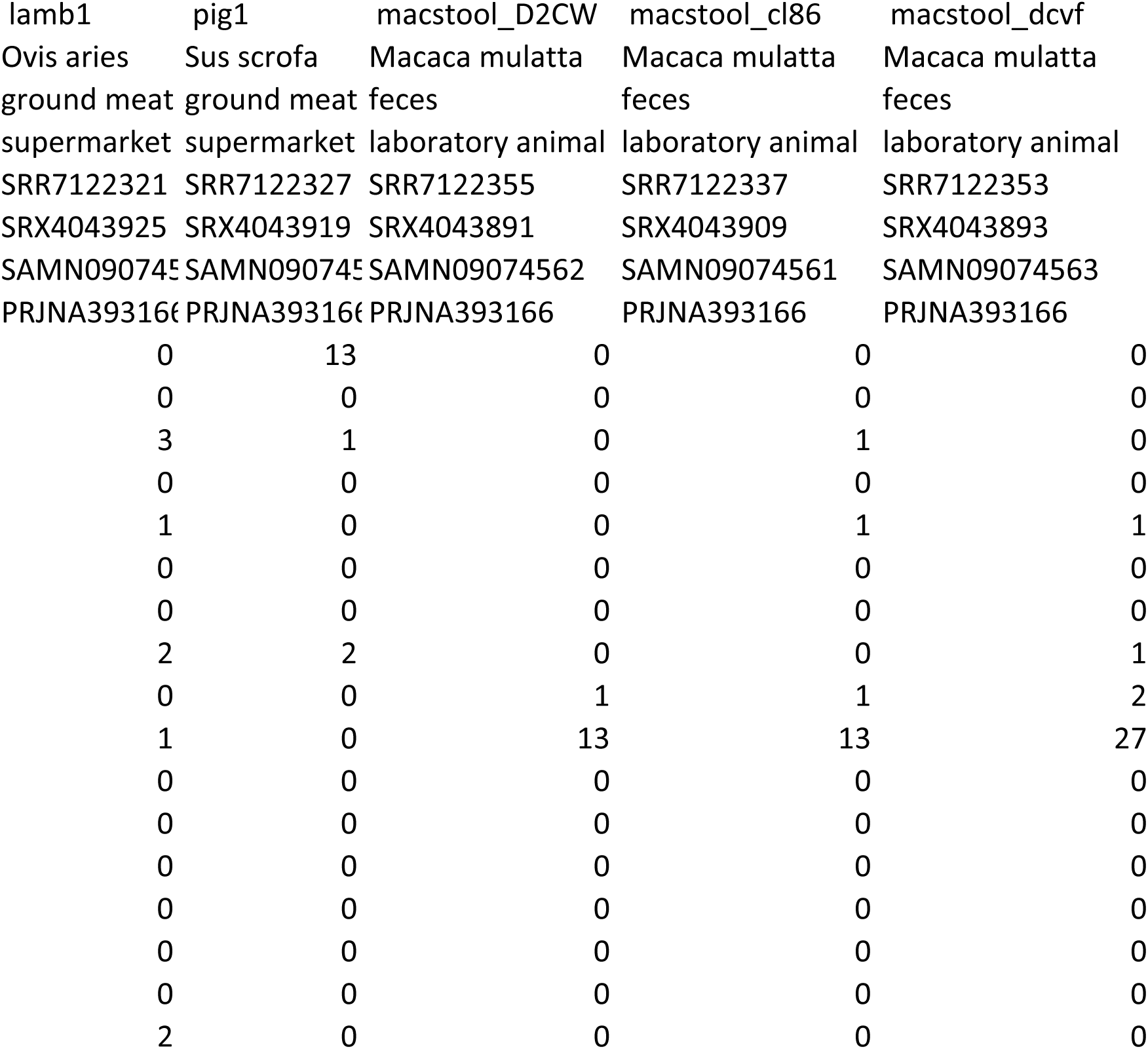
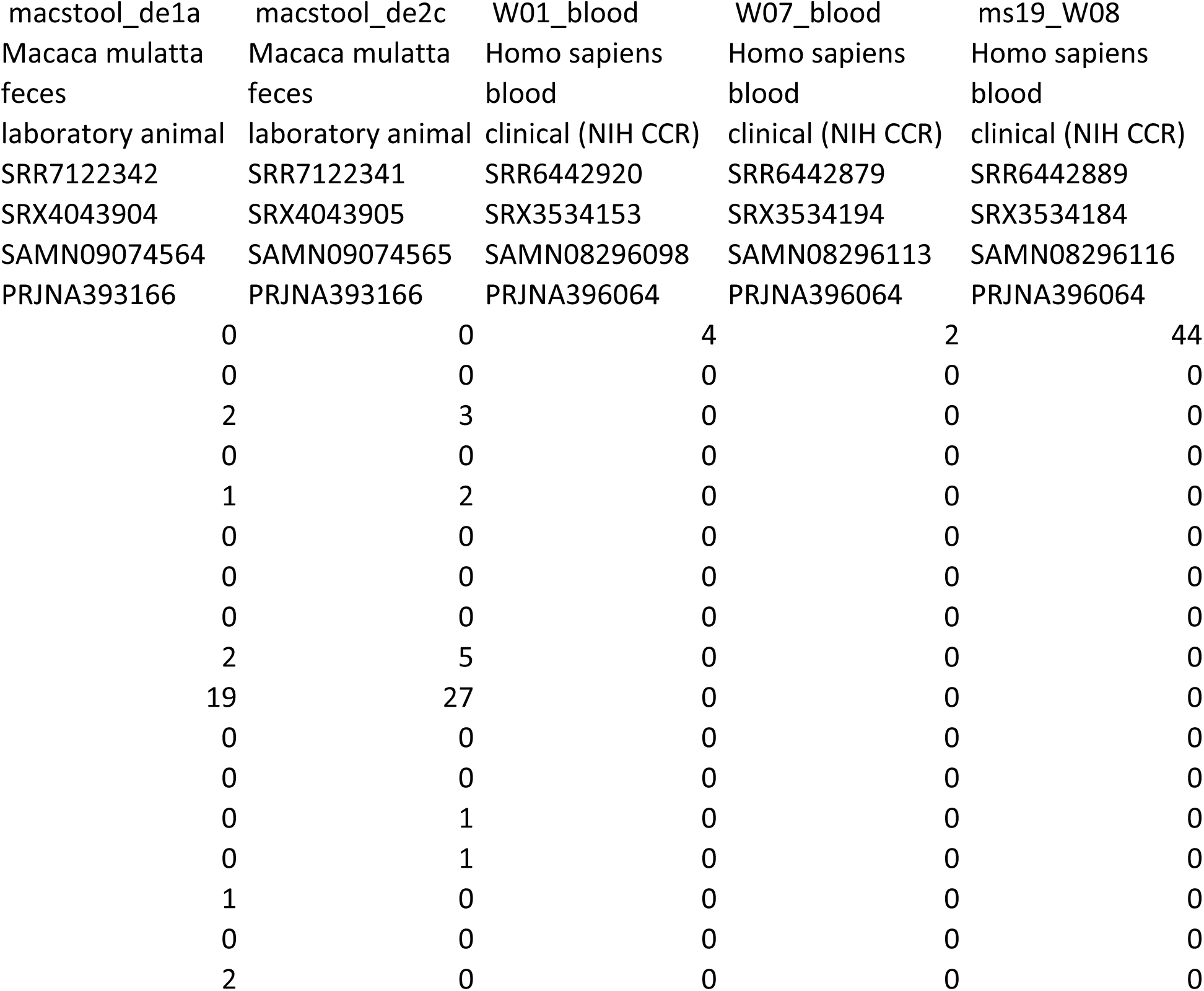
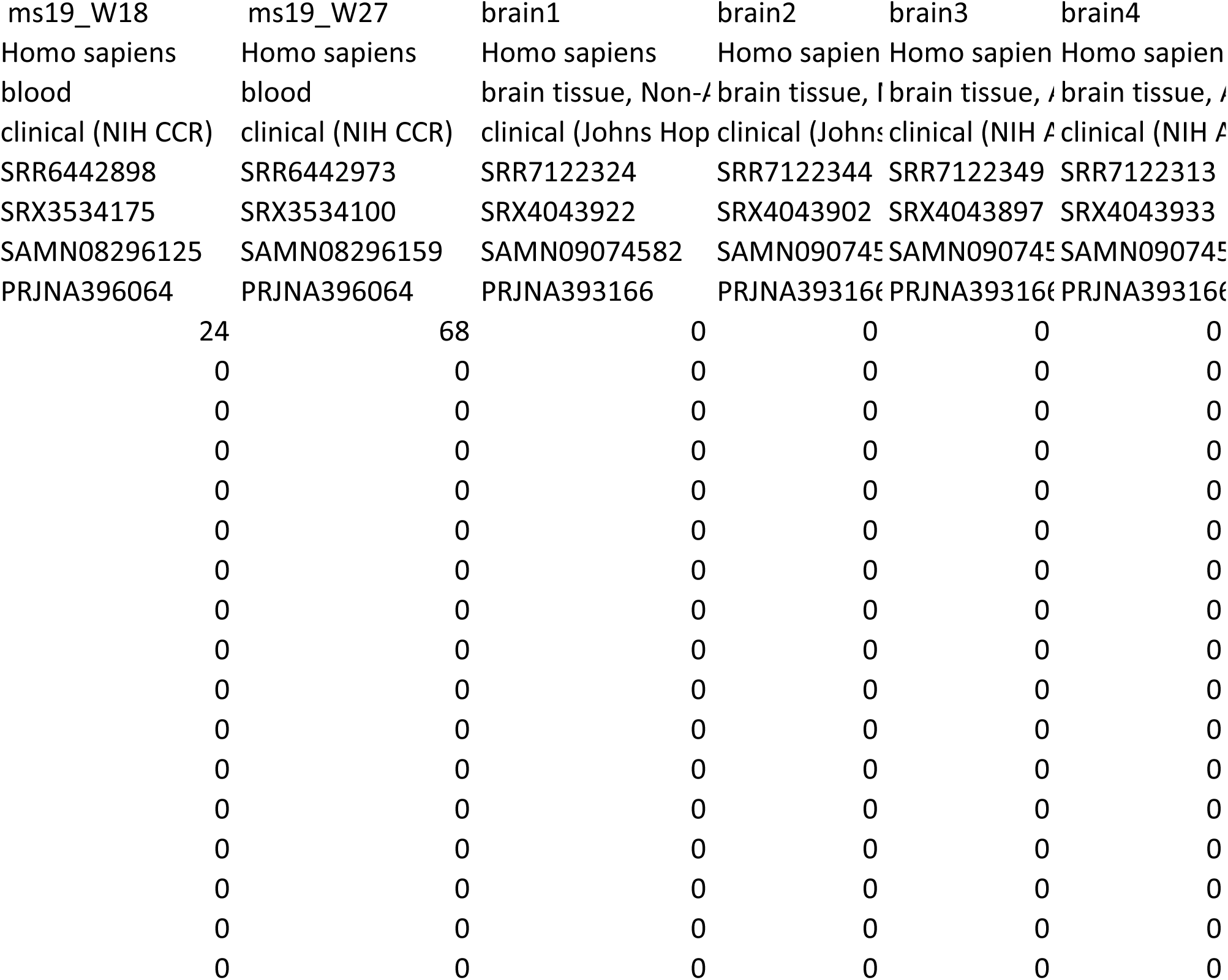
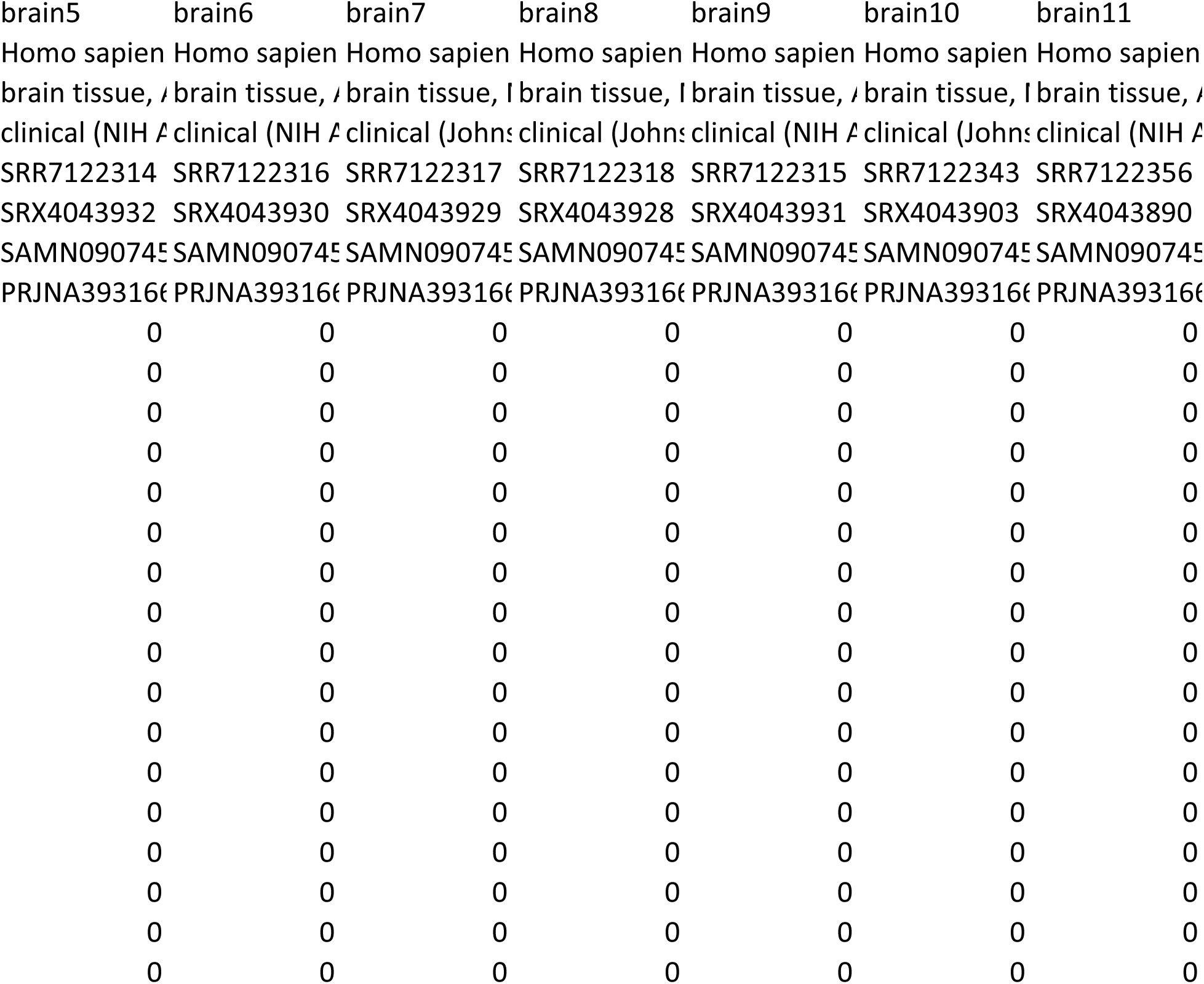
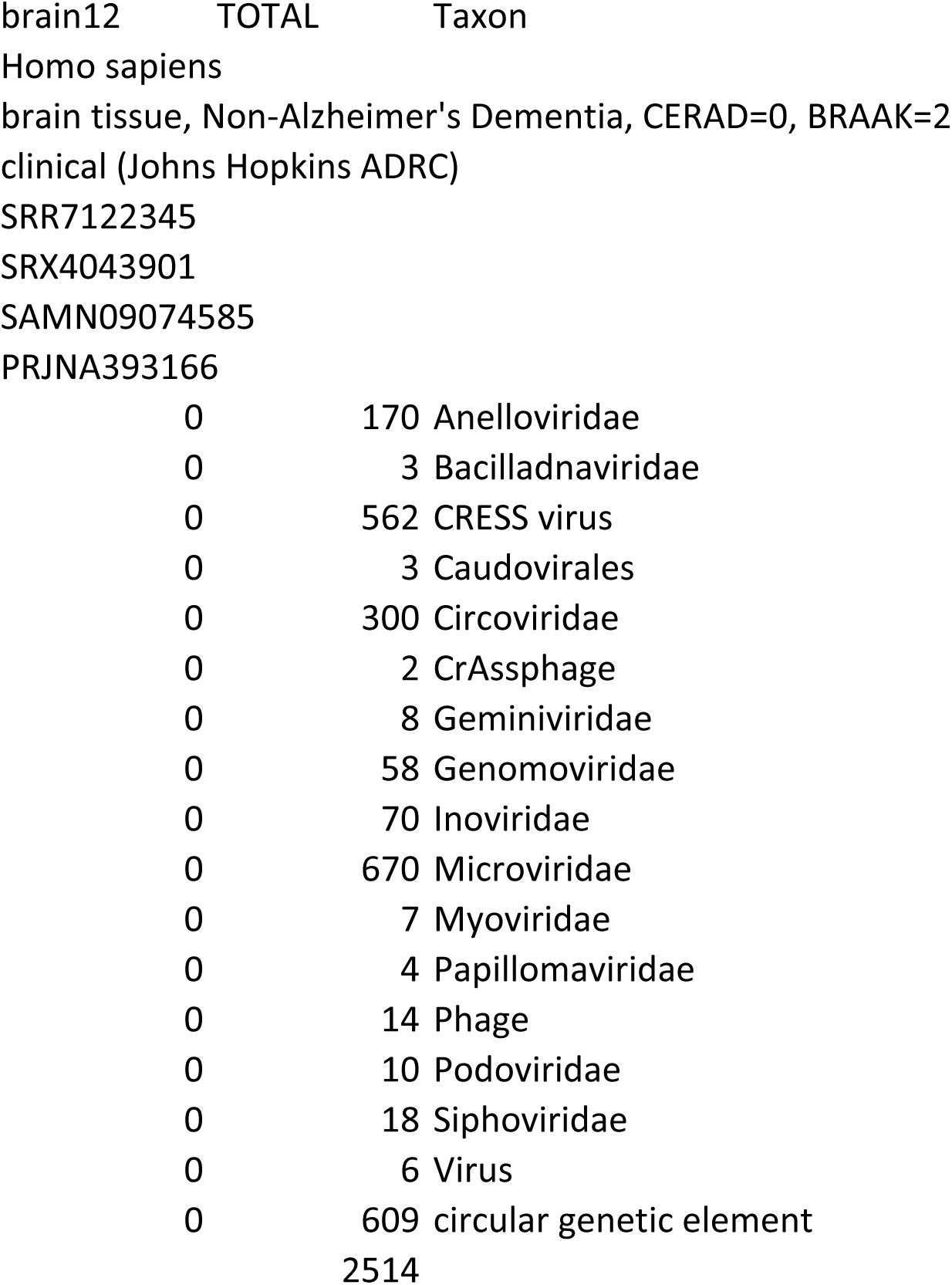

**Table S2.**

**Microviridae sp. isolate ctgd198****MH552479**
| <u>Illumina Library</u> | <u>SRR #</u> | <u>Illumina sequencing run #</u> | <u>Contig Coverage Depth</u> |
| --- | --- | --- | --- |
| 10mus | SRR7122376 | 18 | 97.2 |
| bison1 | SRR7122319 | 4 | 50.5 |
| haddock1 | SRR5832142 | 12 | 18.7 |
| rainbowtrout1 | SRR7122364 | 14 | 37.9 |

**Inoviridae sp. isolate ctda6****MH552488**
| <u>Illumina Library</u> | <u>SRR #</u> | <u>Illumina sequencing run #</u> | <u>Contig Coverage Depth</u> |
| --- | --- | --- | --- |
| abalone1 | SRR7122363 | 12 | 21.8 |
| W01_blood | SRR6442920 | 19 | 62.4 |
| ms19_W08 | SRR6442889 | 19 | 3.5 |
| ms19_W18 | SRR6442898 | 19 | 2.5 |
| ms19_W27 | SRR6442973 | 19 | 3.4 |

**CRESS virus sp. isolate ctbd281****MH617267**
| <u>Illumina Library</u> | <u>SRR #</u> | <u>Illumina sequencing run #</u> | <u>Contig Coverage Depth</u> |
| --- | --- | --- | --- |
| abalone1 | SRR7122363 | 12 | 8.1 |
| bison2 | SRR7122320 | 4 | 84.9 |
| haddock1 | SRR5832142 | 12 | 73.9 |
| squid1 | SRR7122358 | 12 | 4.6 |

**CRESS virus sp. isolate cthf761****MH552486**
| <u>Illumina Library</u> | <u>SRR #</u> | <u>Illumina sequencing run #</u> | <u>Contig Coverage Depth</u> |
| --- | --- | --- | --- |
| 10mus | SRR7122376 | 18 | 65.1 |
| apo1 | SRR7122347 | 18 | 115.7 |
| redsnapper1 | SRR7122366 | 3 | 31.4 |

**Circular genetic element sp. isolate ctbf001****MH648779**
| <u>Illumina Library</u> | <u>SRR #</u> | <u>Illumina sequencing run #</u> | <u>Contig Coverage Depth</u> |
| --- | --- | --- | --- |
| beef_bd3 | SRR7122336 | 3 | 28.7 |
| raccoonskin1 | SRR7122367 | 8 | 10.1 |

**Circular genetic element sp. isolate ctbi451** **MK012536**
| <u>Illumina Library</u> | <u>SRR #</u> | <u>Illumina sequencing run #</u> | <u>Contig Coverage Depth</u> |
| --- | --- | --- | --- |
| 10mus | SRR7122376 | 18 | 65.1 |
| apo1 | SRR7122347 | 18 | 115.7 |
| beef_bd3 | SRR7122336 | 3 | 11.1 |
| bison1 | SRR7122319 | 4 | 11.3 |
| bison2 | SRR7122320 | 4 | 5 |
| lamb1 | SRR7122321 | 4 | 9.4 |
| haddock1 | SRR5832142 | 12 | 8.9 |
| redsnapper1 | SRR7122366 | 3 | 31.4 |
| turkey | SRR7122322 | 4 | 6 |

## References

Agranovsky, A. A., Lesemann, D. E., Maiss, E., Hull, R. & Atabekov, J. G. 1995. “Rattlesnake” structure of a filamentous plant Rna virus built of two capsid proteins. Proc Natl Acad Sci U S A, 92, 2470–3.

Altschul, S. F., Gish, W., Miller, W., Myers, E. W. & Lipman, D. J. 1990. Basic local alignment search tool. J Mol Biol, 215, 403–10.

Asplund, M., Kjartansdottir, K. R., Mollerup, S., Vinner, L., Fridholm, H., Herrera, J. A. R., Friis-Nielsen, J., Hansen, T. A., Jensen, R. H., Nielsen, I. B., Richter, S. R., Rey-Iglesia, A., Matey-Hernandez, M. L., Alquezar-Planas, D. E., Olsen, P. V. S., Sicheritz-Ponten, T., Willerslev, E., Lund, O., Brunak, S., Mourier, T., Nielsen, L. P., Izarzugaza, J. M. G. & Hansen, A. J. 2019. Contaminating viral sequences in high-throughput sequencing viromics: a linkage study of 700 sequencing libraries. Clin Microbiol Infect.

Bedell, M. A., Hudson, J. B., Golub, T. R., Turyk, M. E., Hosken, M., Wilbanks, G. D. & Laimins, L. A. 1991. Amplification of human papillomavirus genomes in vitro is dependent on epithelial differentiation. J Virol, 65, 2254–60.

Bin Jang, H., Bolduc, B., Zablocki, O., Kuhn, J. H., Roux, S., Adriaenssens, E. M., Brister, J. R., Kropinski, A. M., Krupovic, M., Lavigne, R., Turner, D. & Sullivan, M. B. 2019. Taxonomic assignment of uncultivated prokaryotic virus genomes is enabled by gene-sharing networks. Nat Biotechnol, 37, 632–639.

Blinkova, O., Victoria, J., Li, Y., Keele, B. F., Sanz, C., Ndjango, J. B., Peeters, M., Travis, D., Lonsdorf, E. V., Wilson, M. L., Pusey, A. E., Hahn, B. H. & Delwart, E. L. 2010. Novel circular Dna viruses in stool samples of wild-living chimpanzees. J Gen Virol, 91, 74–86.

Bolduc, B., Jang, H. B., Doulcier, G., You, Z. Q., Roux, S. & Sullivan, M. B. 2017. vConTACT: an iVirus tool to classify double-stranded Dna viruses that infect Archaea and Bacteria. PeerJ, 5, e3243.

Bottcher, B., Unseld, S., Ceulemans, H., Russell, R. B. & Jeske, H. 2004. Geminate structures of African cassava mosaic virus. J Virol, 78, 6758–65.

Brister, J. R., Ako-Adjei, D., Bao, Y. & Blinkova, O. 2015. Ncbi viral genomes resource. Nucleic Acids Res, 43, D571–7.

Buck, C. B., Pastrana, D. V., Lowy, D. R. & Schiller, J. T. 2004. Efficient intracellular assembly of papillomaviral vectors. J Virol, 78, 751–7.

Buck, C. B., Van Doorslaer, K., Peretti, A., Geoghegan, E. M., Tisza, M. J., An, P., Katz, J. P., Pipas, J. M., Mcbride, A. A., Camus, A. C., Mcdermott, A. J., Dill, J. A., Delwart, E., Ng, T. F., Farkas, K., Austin, C., Kraberger, S., Davison, W., Pastrana, D. V. & Varsani, A. 2016. The Ancient Evolutionary History of Polyomaviruses. PloS Pathog, 12, e1005574.

Burley, S. K., Berman, H. M., Kleywegt, G. J., Markley, J. L., Nakamura, H. & Velankar, S. 2017. Protein Data Bank (Pdb): The Single Global Macromolecular Structure Archive. Methods Mol Biol, 1607, 627–641.

Chalkias, S., Gorham, J. M., Mazaika, E., Parfenov, M., Dang, X., Depalma, S., Mckean, D., Seidman, C. E., Seidman, J. G. & Koralnik, I. J. 2018. ViroFind: A novel target-enrichment deep-sequencing platform reveals a complex Jc virus population in the brain of Pml patients. PloS One, 13, e0186945.

Chandonia, J. M., Fox, N. K. & Brenner, S. E. 2019. Scope: classification of large macromolecular structures in the structural classification of proteins-extended database. Nucleic Acids Res, 47, D475–D481.

Chen, J., Tsai, V., Parker, W. E., Aronica, E., Baybis, M. & Crino, P. B. 2012. Detection of human papillomavirus in human focal cortical dysplasia type Iib. Ann Neurol, 72, 881–92.

Coras, R., Korn, K., Bien, C. G., Kalbhenn, T., Rossler, K., Kobow, K., Giedl, J., Fleckenstein, B. & Blumcke, I. 2015. No evidence for human papillomavirus infection in focal cortical dysplasia Iib. Ann Neurol, 77, 312–9.

Dayaram, A., Galatowitsch, M. L., Arguello-Astorga, G. R., Van Bysterveldt, K., Kraberger, S., Stainton, D., Harding, J. S., Roumagnac, P., Martin, D. P., Lefeuvre, P. & Varsani, A. 2016. Diverse circular replication-associated protein encoding viruses circulating in invertebrates within a lake ecosystem. Infect Genet Evol, 39, 304–316.

Dayaram, A., Goldstien, S., Arguello-Astorga, G. R., Zawar-Reza, P., Gomez, C., Harding, J. S. & Varsani, A. 2015. Diverse small circular Dna viruses circulating amongst estuarine molluscs. Infect Genet Evol, 31, 284–95.

Díez-Villaseñor, C. & Rodriguez-Valera, F. 2019. Crispr analysis suggests that small circular single-stranded Dna smacoviruses infect Archaea instead of humans. Nature Communications, 10, 294.

Eimer, W. A., Vijaya Kumar, D. K., Navalpur Shanmugam, N. K., Rodriguez, A. S., Mitchell, T., Washicosky, K. J., Gyorgy, B., Breakefield, X. O., Tanzi, R. E. & Moir, R. D. 2018. Alzheimer’s Disease-Associated beta-Amyloid Is Rapidly Seeded by Herpesviridae to Protect against Brain Infection. Neuron, 99, 56–63 e3.

El-Gebali, S., Mistry, J., Bateman, A., Eddy, S. R., Luciani, A., Potter, S. C., Qureshi, M., Richardson, L. J., Salazar, G. A., Smart, A., Sonnhammer, E. L. L., Hirsh, L., Paladin, L., Piovesan, D., Tosatto, S. C. E. & Finn, R. D. 2019. The Pfam protein families database in 2019. Nucleic Acids Res, 47, D427–D432.

Emerson, J. B., Roux, S., Brum, J. R., Bolduc, B., Woodcroft, B. J., Jang, H. B., Singleton, C. M., Solden, L. M., Naas, A. E., Boyd, J. A., Hodgkins, S. B., Wilson, R. M., Trubl, G., Li, C., Frolking, S., Pope, P. B., Wrighton, K. C., Crill, P. M., Chanton, J. P., Saleska, S. R., Tyson, G. W., Rich, V. I. & Sullivan, M. B. 2018. Host-linked soil viral ecology along a permafrost thaw gradient. Nat Microbiol, 3, 870–880.

Falero, A., Caballero, A., Ferran, B., Izquierdo, Y., Fando, R. & Campos, J. 2009. Dna binding proteins of the filamentous phages Ctxphi and Vgjphi of Vibrio cholerae. J Bacteriol, 191, 5873–6.

Fontenele, R. S., Lacorte, C., Lamas, N. S., Schmidlin, K., Varsani, A. & Ribeiro, S. G. 2019. Single Stranded Dna Viruses Associated with Capybara Faeces Sampled in Brazil. Viruses, 11.

Fu, L., Niu, B., Zhu, Z., Wu, S. & Li, W. 2012. Cd-Hit: accelerated for clustering the next-generation sequencing data. Bioinformatics, 28, 3150–2.

Geoghegan, E. M., Pastrana, D. V., Schowalter, R. M., Ray, U., Gao, W., Ho, M., Pauly, G. T., Sigano, D. M., Kaynor, C., Cahir-Mcfarland, E., Combaluzier, B., Grimm, J. & Buck, C. B. 2017. Infectious Entry and Neutralization of Pathogenic Jc Polyomaviruses. Cell Rep, 21, 1169–1179.

Gerlt, J. A., Bouvier, J. T., Davidson, D. B., Imker, H. J., Sadkhin, B., Slater, D. R. & Whalen, K. L. 2015. Enzyme Function Initiative-Enzyme Similarity Tool (Efi-Est): A web tool for generating protein sequence similarity networks. Biochim Biophys Acta, 1854, 1019–37.

Gilbert, J. A., Meyer, F., Antonopoulos, D., Balaji, P., Brown, C. T., Brown, C. T., Desai, N., Eisen, J. A., Evers, D., Field, D., Feng, W., Huson, D., Jansson, J., Knight, R., Knight, J., Kolker, E., Konstantindis, K., Kostka, J., Kyrpides, N., Mackelprang, R., Mchardy, A., Quince, C., Raes, J., Sczyrba, A., Shade, A. & Stevens, R. 2010. Meeting report: the terabase metagenomics workshop and the vision of an Earth microbiome project. Stand Genomic Sci, 3, 243–8.

Gregory, A. C., Zayed, A. A., Conceicao-Neto, N., Temperton, B., Bolduc, B., Alberti, A., Ardyna, M., Arkhipova, K., Carmichael, M., Cruaud, C., Dimier, C., Dominguez-Huerta, G., Ferland, J., Kandels, S., Liu, Y., Marec, C., Pesant, S., Picheral, M., Pisarev, S., Poulain, J., Tremblay, J. E., Vik, D., Tara Oceans, C., Babin, M., Bowler, C., Culley, A. I., De Vargas, C., Dutilh, B. E., Iudicone, D., Karp-Boss, L., Roux, S., Sunagawa, S., Wincker, P. & Sullivan, M. B. 2019. Marine Dna Viral Macro- and Microdiversity from Pole to Pole. Cell, 177, 1109–1123 e14.

Greninger, A. L. & Derisi, J. L. 2015. Draft Genome Sequences of Ciliovirus and Brinovirus from San Francisco Wastewater. Genome Announc, 3.

Grose, J. H. & Casjens, S. R. 2014. Understanding the enormous diversity of bacteriophages: the tailed phages that infect the bacterial family Enterobacteriaceae. Virology, 468-470, 421–443.

Grose, J. H., Jensen, G. L., Burnett, S. H. & Breakwell, D. P. 2014. Genomic comparison of 93 Bacillus phages reveals 12 clusters, 14 singletons and remarkable diversity. Bmc Genomics, 15, 855.

Hipp, K., Grimm, C., Jeske, H. & Bottcher, B. 2017. Near-Atomic Resolution Structure of a Plant Geminivirus Determined by Electron Cryomicroscopy. Structure, 25, 1303–1309 e3.

Hodyra-Stefaniak, K., Miernikiewicz, P., Drapala, J., Drab, M., Jonczyk-Matysiak, E., Lecion, D., Kazmierczak, Z., Beta, W., Majewska, J., Harhala, M., Bubak, B., Klopot, A., Gorski, A. & Dabrowska, K. 2015. Mammalian Host-Versus-Phage immune response determines phage fate in vivo. Sci Rep, 5, 14802.

Huang, Y. J., Mao, B., Aramini, J. M. & Montelione, G. T. 2014. Assessment of template-based protein structure predictions in Casp10. Proteins, 82 Suppl 2, 43–56.

Hunt, M., Silva, N. D., Otto, T. D., Parkhill, J., Keane, J. A. & Harris, S. R. 2015. Circlator: automated circularization of genome assemblies using long sequencing reads. Genome Biol, 16, 294.

Iranzo, J., Krupovic, M. & Koonin, E. V. 2017. A network perspective on the virus world. Commun Integr Biol, 10, e1296614.

Itzhaki, R. F., Lathe, R., Balin, B. J., Ball, M. J., Bearer, E. L., Braak, H., Bullido, M. J., Carter, C., Clerici, M., Cosby, S. L., Del Tredici, K., Field, H., Fulop, T., Grassi, C., Griffin, W. S., Haas, J., Hudson, A. P., Kamer, A. R., Kell, D. B., Licastro, F., Letenneur, L., Lovheim, H., Mancuso, R., Miklossy, J., Otth, C., Palamara, A. T., Perry, G., Preston, C., Pretorius, E., Strandberg, T., Tabet, N., Taylor-Robinson, S. D. & Whittum-Hudson, J. A. 2016. Microbes and Alzheimer’s Disease. J Alzheimers Dis, 51, 979–84.

Katoh, K. & Standley, D. M. 2013. Mafft multiple sequence alignment software version 7: improvements in performance and usability. Mol Biol Evol, 30, 772–80.

Kauffman, K. M., Hussain, F. A., Yang, J., Arevalo, P., Brown, J. M., Chang, W. K., Vaninsberghe, D., Elsherbini, J., Sharma, R. S., Cutler, M. B., Kelly, L. & Polz, M. F. 2018. A major lineage of non-tailed dsdna viruses as unrecognized killers of marine bacteria. Nature, 554, 118–122.

Kazlauskas, D., Dayaram, A., Kraberger, S., Goldstien, S., Varsani, A. & Krupovic, M. 2017. Evolutionary history of ssdna bacilladnaviruses features horizontal acquisition of the capsid gene from ssrna nodaviruses. Virology, 504, 114–121.

Kazlauskas, D., Varsani, A., Koonin, E. V. & Krupovic, M. 2019. Multiple origins of prokaryotic and eukaryotic single-stranded Dna viruses from bacterial and archaeal plasmids. Nat Commun, 10, 3425.

Kelley, L. A., Mezulis, S., Yates, C. M., Wass, M. N. & Sternberg, M. J. 2015. The Phyre2 web portal for protein modeling, prediction and analysis. Nat Protoc, 10, 845–58.

Khayat, R., Brunn, N., Speir, J. A., Hardham, J. M., Ankenbauer, R. G., Schneemann, A. & Johnson, J. E. 2011. The 2.3-angstrom structure of porcine circovirus 2. J Virol, 85, 7856–62.

Kim, K. H., Chang, H. W., Nam, Y. D., Roh, S. W., Kim, M. S., Sung, Y., Jeon, C. O., Oh, H. M. & Bae, J. W. 2008. Amplification of uncultured single-stranded Dna viruses from rice paddy soil. Appl Environ Microbiol, 74, 5975–85.

King, A. M. Q., Lefkowitz, E. J., Mushegian, A. R., Adams, M. J., Dutilh, B. E., Gorbalenya, A. E., Harrach, B., Harrison, R. L., Junglen, S., Knowles, N. J., Kropinski, A. M., Krupovic, M., Kuhn, J. H., Nibert, M. L., Rubino, L., Sabanadzovic, S., Sanfacon, H., Siddell, S. G., Simmonds, P., Varsani, A., Zerbini, F. M. & Davison, A. J. 2018. Changes to taxonomy and the International Code of Virus Classification and Nomenclature ratified by the International Committee on Taxonomy of Viruses (2018). Arch Virol.

Koonin, E. V., Dolja, V. V. & Krupovic, M. 2015. Origins and evolution of viruses of eukaryotes: The ultimate modularity. Virology, 479-480, 2–25.

Kraberger, S., Arguello-Astorga, G. R., Greenfield, L. G., Galilee, C., Law, D., Martin, D. P. & Varsani, A. 2015. Characterisation of a diverse range of circular replication-associated protein encoding Dna viruses recovered from a sewage treatment oxidation pond. Infect Genet Evol, 31, 73–86.

Kraberger, S., Schmidlin, K., Fontenele, R. S., Walters, M. & Varsani, A. 2019. Unravelling the Single-Stranded Dna Virome of the New Zealand Blackfly. Viruses, 11.

Krishnamurthy, S. R. & Wang, D. 2017. Origins and challenges of viral dark matter. Virus Res, 239, 136–142.

Krupovic, M., Ghabrial, S. A., Jiang, D. & Varsani, A. 2016. Genomoviridae: a new family of widespread single-stranded Dna viruses. Arch Virol, 161, 2633–43.

Krupovic, M. & Koonin, E. V. 2017. Multiple origins of viral capsid proteins from cellular ancestors. Proc Natl Acad Sci U S A, 114, E2401–E2410.

Krupovic, M., Ravantti, J. J. & Bamford, D. H. 2009. Geminiviruses: a tale of a plasmid becoming a virus. Bmc Evol Biol, 9, 112.

Krupovic, M., Zhi, N., Li, J., Hu, G., Koonin, E. V., Wong, S., Shevchenko, S., Zhao, K. & Young, N. S. 2015. Multiple layers of chimerism in a single-stranded Dna virus discovered by deep sequencing. Genome Biol Evol, 7, 993–1001.

Labonte, J. M. & Suttle, C. A. 2013. Previously unknown and highly divergent ssdna viruses populate the oceans. Isme J, 7, 2169–77.

Lefeuvre, P., Martin, D. P., Elena, S. F., Shepherd, D. N., Roumagnac, P. & Varsani, A. 2019. Evolution and ecology of plant viruses. Nat Rev Microbiol.

Letunic, I. & Bork, P. 2019. Interactive Tree Of Life (itol) v4: recent updates and new developments. Nucleic Acids Res, 47, W256–W259.

Lima-Mendez, G., Van Helden, J., Toussaint, A. & Leplae, R. 2008. Reticulate representation of evolutionary and functional relationships between phage genomes. Mol Biol Evol, 25, 762–77.

Lloyd-Price, J., Mahurkar, A., Rahnavard, G., Crabtree, J., Orvis, J., Hall, A. B., Brady, A., Creasy, H. H., Mccracken, C., Giglio, M. G., Mcdonald, D., Franzosa, E. A., Knight, R., White, O. & Huttenhower, C. 2017. Strains, functions and dynamics in the expanded Human Microbiome Project. Nature, 550, 61–66.

Makino, D. L., Larson, S. B. & Mcpherson, A. 2013. The crystallographic structure of Panicum Mosaic Virus (Pmv). J Struct Biol, 181, 37–52.

Marchler-Bauer, A. & Bryant, S. H. 2004. Cd-Search: protein domain annotations on the fly. Nucleic Acids Res, 32, W327–31.

Marchler-Bauer, A., Derbyshire, M. K., Gonzales, N. R., Lu, S., Chitsaz, F., Geer, L. Y., Geer, R. C., He, J., Gwadz, M., Hurwitz, D. I., Lanczycki, C. J., Lu, F., Marchler, G. H., Song, J. S., Thanki, N., Wang, Z., Yamashita, R. A., Zhang, D., Zheng, C. & Bryant, S. H. 2015. Cdd: Ncbi’s conserved domain database. Nucleic Acids Res, 43, D222–6.

Meier, A. & Soding, J. 2015. Automatic Prediction of Protein 3D Structures by Probabilistic Multi-template Homology Modeling. PloS Comput Biol, 11, e1004343.

Nguyen, L. T., Schmidt, H. A., Von Haeseler, A. & Minh, B. Q. 2015. Iq-Tree: a fast and effective stochastic algorithm for estimating maximum-likelihood phylogenies. Mol Biol Evol, 32, 268–74.

O’leary, N. A., Wright, M. W., Brister, J. R., Ciufo, S., Haddad, D., Mcveigh, R., Rajput, B., Robbertse, B., Smith-White, B., Ako-Adjei, D., Astashyn, A., Badretdin, A., Bao, Y., Blinkova, O., Brover, V., Chetvernin, V., Choi, J., Cox, E., Ermolaeva, O., Farrell, C. M., Goldfarb, T., Gupta, T., Haft, D., Hatcher, E., Hlavina, W., Joardar, V. S., Kodali, V. K., Li, W., Maglott, D., Masterson, P., Mcgarvey, K. M., Murphy, M. R., O’neill, K., Pujar, S., Rangwala, S. H., Rausch, D., Riddick, L. D., Schoch, C., Shkeda, A., Storz, S. S., Sun, H., Thibaud-Nissen, F., Tolstoy, I., Tully, R. E., Vatsan, A. R., Wallin, C., Webb, D., Wu, W., Landrum, M. J., Kimchi, A., Tatusova, T., Dicuccio, M., Kitts, P., Murphy, T. D. & Pruitt, K. D. 2016. Reference sequence (RefSeq) database at Ncbi: current status, taxonomic expansion, and functional annotation. Nucleic Acids Res, 44, D733–45.

Oh, J., Byrd, A. L., Deming, C., Conlan, S., Kong, H. H., Segre, J. A. & Program, N. C. S. 2014. Biogeography and individuality shape function in the human skin metagenome. Nature, 514, 59–64.

Paez-Espino, D., Eloe-Fadrosh, E. A., Pavlopoulos, G. A., Thomas, A. D., Huntemann, M., Mikhailova, N., Rubin, E., Ivanova, N. N. & Kyrpides, N. C. 2016. Uncovering Earth’s virome. Nature, 536, 425–30.

Paez-Espino, D., Roux, S., Chen, I. A., Palaniappan, K., Ratner, A., Chu, K., Huntemann, M., Reddy, T. B. K., Pons, J. C., Llabres, M., Eloe-Fadrosh, E. A., Ivanova, N. N. & Kyrpides, N. C. 2019. Img/Vr v.2.0: an integrated data management and analysis system for cultivated and environmental viral genomes. Nucleic Acids Res, 47, D678–D686.

Pastrana, D. V., Peretti, A., Welch, N. L., Borgogna, C., Olivero, C., Badolato, R., Notarangelo, L. D., Gariglio, M., Fitzgerald, P. C., Mcintosh, C. E., Reeves, J., Starrett, G. J., Bliskovsky, V., Velez, D., Brownell, I., Yarchoan, R., Wyvill, K. M., Uldrick, T. S., Maldarelli, F., Lisco, A., Sereti, I., Gonzalez, C. M., Androphy, E. J., Mcbride, A. A., Van Doorslaer, K., Garcia, F., Dvoretzky, I., Liu, J. S., Han, J., Murphy, P. M., Mcdermott, D. H. & Buck, C. B. 2018. Metagenomic Discovery of 83 New Human Papillomavirus Types in Patients with Immunodeficiency. mSphere, 3.

Pei, J. & Grishin, N. V. 2014. Promals3D: multiple protein sequence alignment enhanced with evolutionary and three-dimensional structural information. Methods Mol Biol, 1079, 263–71.

Peretti, A., Fitzgerald, P. C., Bliskovsky, V., Buck, C. B. & Pastrana, D. V. 2015. Hamburger polyomaviruses. J Gen Virol, 96, 833–9.

Pope, W. H., Bowman, C. A., Russell, D. A., Jacobs-Sera, D., Asai, D. J., Cresawn, S. G., Jacobs, W. R., Hendrix, R. W., Lawrence, J. G., Hatfull, G. F., Science Education Alliance Phage Hunters Advancing, G., Evolutionary, S., Phage Hunters Integrating, R., Education & Mycobacterial Genetics, C. 2015. Whole genome comparison of a large collection of mycobacteriophages reveals a continuum of phage genetic diversity. Elife, 4, e06416.

Quaiser, A., Krupovic, M., Dufresne, A., Francez, A. J. & Roux, S. 2016. Diversity and comparative genomics of chimeric viruses in Sphagnum-dominated peatlands. Virus Evol, 2, vew025.

Remmert, M., Biegert, A., Hauser, A. & Söding, J. 2011. Hhblits: lightning-fast iterative protein sequence searching by Hmm-Hmm alignment. Nat Methods, 9, 173–5.

Rohwer, F. & Edwards, R. 2002. The Phage Proteomic Tree: a genome-based taxonomy for phage. J Bacteriol, 184, 4529–35.

Rosario, K., Breitbart, M., Harrach, B., Segales, J., Delwart, E., Biagini, P. & Varsani, A. 2017. Revisiting the taxonomy of the family Circoviridae: establishment of the genus Cyclovirus and removal of the genus Gyrovirus. Arch Virol, 162, 1447–1463.

Rosario, K., Mettel, K. A., Benner, B. E., Johnson, R., Scott, C., Yusseff-Vanegas, S. Z., Baker, C. C. M., Cassill, D. L., Storer, C., Varsani, A. & Breitbart, M. 2018. Virus discovery in all three major lineages of terrestrial arthropods highlights the diversity of single-stranded Dna viruses associated with invertebrates. PeerJ, 6, e5761.

Roux, S., Adriaenssens, E. M., Dutilh, B. E., Koonin, E. V., Kropinski, A. M., Krupovic, M., Kuhn, J. H., Lavigne, R., Brister, J. R., Varsani, A., Amid, C., Aziz, R. K., Bordenstein, S. R., Bork, P., Breitbart, M., Cochrane, G. R., Daly, R. A., Desnues, C., Duhaime, M. B., Emerson, J. B., Enault, F., Fuhrman, J. A., Hingamp, P., Hugenholtz, P., Hurwitz, B. L., Ivanova, N. N., Labonte, J. M., Lee, K. B., Malmstrom, R. R., Martinez-Garcia, M., Mizrachi, I. K., Ogata, H., Paez-Espino, D., Petit, M. A., Putonti, C., Rattei, T., Reyes, A., Rodriguez-Valera, F., Rosario, K., Schriml, L., Schulz, F., Steward, G. F., Sullivan, M. B., Sunagawa, S., Suttle, C. A., Temperton, B., Tringe, S. G., Thurber, R. V., Webster, N. S., Whiteson, K. L., Wilhelm, S. W., Wommack, K. E., Woyke, T., Wrighton, K. C., Yilmaz, P., Yoshida, T., Young, M. J., Yutin, N., Allen, L. Z., Kyrpides, N. C. & Eloe-Fadrosh, E. A. 2019a. Minimum Information about an Uncultivated Virus Genome (MiuviG). Nature Biotechnology, 37, 29–37.

Roux, S., Enault, F., Bronner, G., Vaulot, D., Forterre, P. & Krupovic, M. 2013. Chimeric viruses blur the borders between the major groups of eukaryotic single-stranded Dna viruses. Nat Commun, 4, 2700.

Roux, S., Enault, F., Hurwitz, B. L. & Sullivan, M. B. 2015. VirSorter: mining viral signal from microbial genomic data. PeerJ, 3, e985.

Roux, S., Krupovic, M., Daly, R. A., Borges, A. L., Nayfach, S., Schulz, F., Sharrar, A., Matheus Carnevali, P. B., Cheng, J. F., Ivanova, N. N., Bondy-Denomy, J., Wrighton, K. C., Woyke, T., Visel, A., Kyrpides, N. C. & Eloe-Fadrosh, E. A. 2019b. Cryptic inoviruses revealed as pervasive in bacteria and archaea across Earth’s biomes. Nat Microbiol.

Schulz, F., Yutin, N., Ivanova, N. N., Ortega, D. R., Lee, T. K., Vierheilig, J., Daims, H., Horn, M., Wagner, M., Jensen, G. J., Kyrpides, N. C., Koonin, E. V. & Woyke, T. 2017. Giant viruses with an expanded complement of translation system components. Science, 356, 82–85.

Seguritan, V., Alves, N., Jr., Arnoult, M., Raymond, A., Lorimer, D., Burgin, A. B., Jr., Salamon, P. & Segall, A. M. 2012. Artificial neural networks trained to detect viral and phage structural proteins. PloS Comput Biol, 8, e1002657.

Shi, M., Lin, X. D., Tian, J. H., Chen, L. J., Chen, X., Li, C. X., Qin, X. C., Li, J., Cao, J. P., Eden, J. S., Buchmann, J., Wang, W., Xu, J., Holmes, E. C. & Zhang, Y. Z. 2016. Redefining the invertebrate Rna virosphere. Nature.

Simmonds, P., Adams, M. J., Benko, M., Breitbart, M., Brister, J. R., Carstens, E. B., Davison, A. J., Delwart, E., Gorbalenya, A. E., Harrach, B., Hull, R., King, A. M., Koonin, E. V., Krupovic, M., Kuhn, J. H., Lefkowitz, E. J., Nibert, M. L., Orton, R., Roossinck, M. J., Sabanadzovic, S., Sullivan, M. B., Suttle, C. A., Tesh, R. B., Van Der Vlugt, R. A., Varsani, A. & Zerbini, F. M. 2017. Consensus statement: Virus taxonomy in the age of metagenomics. Nat Rev Microbiol, 15, 161–168.

Steel, O., Kraberger, S., Sikorski, A., Young, L. M., Catchpole, R. J., Stevens, A. J., Ladley, J. J., Coray, D. S., Stainton, D., Dayaram, A., Julian, L., Van Bysterveldt, K. & Varsani, A. 2016. Circular replication-associated protein encoding Dna viruses identified in the faecal matter of various animals in New Zealand. Infect Genet Evol, 43, 151–64.

Su, G., Morris, J. H., Demchak, B. & Bader, G. D. 2014. Biological network exploration with Cytoscape 3. Curr Protoc Bioinformatics, 47, 8.13.1–24.

Sullivan, M. B. 2015. Viromes, not gene markers, for studying double-stranded Dna virus communities. J Virol, 89, 2459–61.

Tomaru, Y., Takao, Y., Suzuki, H., Nagumo, T., Koike, K. & Nagasaki, K. 2011. Isolation and characterization of a single-stranded Dna virus infecting Chaetoceros lorenzianus Grunow. Appl Environ Microbiol, 77, 5285–93.

Turnbaugh, P. J., Ley, R. E., Hamady, M., Fraser-Liggett, C. M., Knight, R. & Gordon, J. I. 2007. The human microbiome project. Nature, 449, 804–10.

Uniprot, C. 2019. UniProt: a worldwide hub of protein knowledge. Nucleic Acids Res, 47, D506–D515.

Varsani, A. & Krupovic, M. 2018. Smacoviridae: a new family of animal-associated single-stranded Dna viruses. Arch Virol, 163, 2005–2015.

Victoria, J. G., Kapoor, A., Li, L., Blinkova, O., Slikas, B., Wang, C., Naeem, A., Zaidi, S. & Delwart, E. 2009. Metagenomic analyses of viruses in stool samples from children with acute flaccid paralysis. J Virol, 83, 4642–51.

Waldor, M. K. & Mekalanos, J. J. 1996. Lysogenic conversion by a filamentous phage encoding cholera toxin. Science, 272, 1910–4.

Wick, R. R., Schultz, M. B., Zobel, J. & Holt, K. E. 2015. Bandage: interactive visualization of de novo genome assemblies. Bioinformatics, 31, 3350–2.

Zerbini, F. M., Briddon, R. W., Idris, A., Martin, D. P., Moriones, E., Navas-Castillo, J., Rivera-Bustamante, R., Roumagnac, P., Varsani, A. & Ictv Report, C. 2017. Ictv Virus Taxonomy Profile: Geminiviridae. J Gen Virol, 98, 131–133.

Zhang, W., Olson, N. H., Baker, T. S., Faulkner, L., Agbandje-Mckenna, M., Boulton, M. I., Davies, J. W. & Mckenna, R. 2001. Structure of the Maize streak virus geminate particle. Virology, 279, 471–7.

Zhao, L., Rosario, K., Breitbart, M. & Duffy, S. 2019. Eukaryotic Circular Rep-Encoding Single-Stranded Dna (Cress Dna) Viruses: Ubiquitous Viruses With Small Genomes and a Diverse Host Range. Adv Virus Res, 103, 71–133.

Zimmermann, L., Stephens, A., Nam, S. Z., Rau, D., Kubler, J., Lozajic, M., Gabler, F., Soding, J., Lupas, A. N. & Alva, V. 2018. A Completely Reimplemented Mpi Bioinformatics Toolkit with a New Hhpred Server at its Core. J Mol Biol, 430, 2237–2243.

